# Hydration and hydrolysis define antibiotic resistance conferred by macrolide esterases

**DOI:** 10.64898/2026.03.24.713787

**Authors:** Emma T. R. Kelly, Iryna Myziuk, Mark Z. Hemmings, Zahi Mulla, Jonathan Blanchet, Antonio Ruzzini, Albert M. Berghuis

## Abstract

Macrolides are an antibiotic class widely used in both human and veterinary medicine, and function by interfering with protein synthesis. Regrettably, numerous strategies for evading the antibiotic properties of macrolides have been found in bacteria, including enzyme-mediated inactivation. These mechanisms are now widely disseminated among pathogenic, animal-associated and environmental bacteria making them a One Health issue. Macrolide esterases, which hydrolyze the macrolactone’s ester bond, confer one such resistance mechanism. Two types of macrolide esterases have thus far been identified, the well-studied erythromycin esterases and the recently discovered Est-type enzymes that belong to the α/β-hydrolase superfamily. We present detailed structure-function studies for four diverse Est type esterases: which only share 44-66% sequence identity (EstT_Sf_, EstT_St_, EstT_Bc_, and EstX_Ec_). In addition to resistance profiling and substrate specificity studies, we present structures for all four enzymes, including structures for EstT_Bc_ and EstX_Ec_ in complex with tylosin and tylvalosin macrolides, post hydrolysis. Complementing the data with mutational and kinetic studies allowed for a detailed analysis of the structural basis for macrolide-enzyme interactions. Combined the data suggest that promiscuous binding and imprecise positioning, mediated by a water-cage, dictate substrate specificity for Est-type macrolide resistance enzymes. These insights may prove beneficial for next-generation antibiotic development.

## Introduction

Macrolides are a class of antibiotics that, since the Golden Age of Antibiotics, have been broadly applied in both clinical and agricultural settings (1). The class represents tens of billions of dollars within the antibiotic global market (2). These antibiotics are used to treat a wide range of bacterial infections. Although they show the greatest activity against Gram-positive bacterial infections, both natural products and semi-synthetics are administered en masse to manage disease and to treat respiratory infections caused by cell wall-free and Gram-negative veterinary pathogens that disrupt food production (1, 3, 4). Macrolides are an actinomycete bioderived product, characterized by a macrocyclic lactone (of varying sizes) that is substituted with at least one deoxy sugar via glycosidic linkage (Figure 1S) (1). Macrolides with 14-, 15-, and 16-membered rings function by targeting the bacterial 50S ribosomal subunit where they bind to the highly conserved nascent peptide exit tunnel, which blocks the lumen and obstructs polypeptide elongation (4, 5). This effect can either be bactericidal or bacteriostatic depending on the macrolide in question.

**Figure 1.**
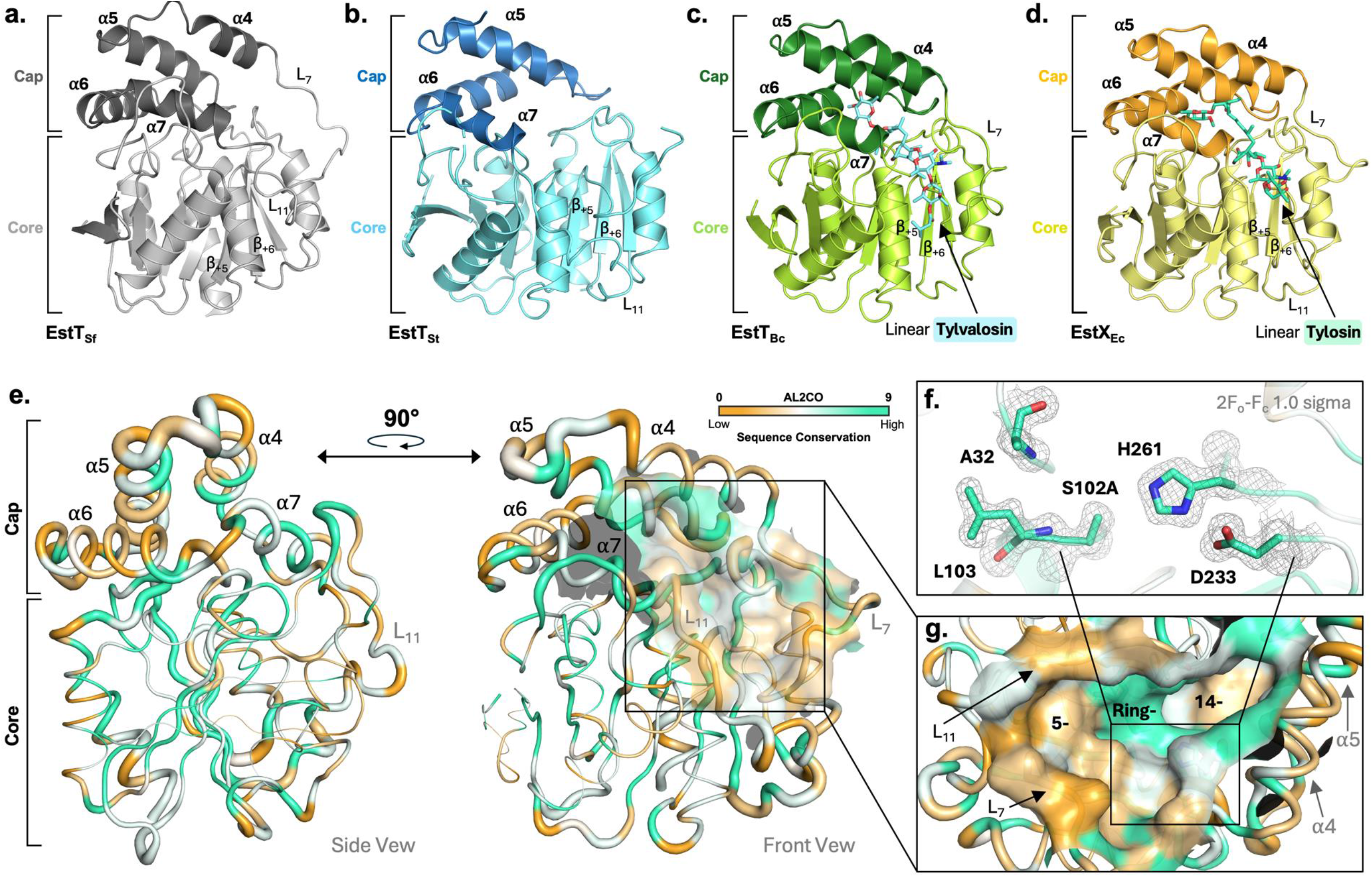
X-ray crystal structures of α/β-hydrolase superfamily macrolide esterase members: **a.** EstT_Sf_ (grey), **b**. EstT_St_ (blue), **c**. EstT_Bc_ (green) in complex with linear tylvalosin (aquamarine), and **d**. EstX_Ec_ (yellow) in complex with linear tylosin (greencyan). α/β-hydrolase core domains are shown in light colours, immobile cap domains are down in dark. **e**. Structure and sequence conservation of the four Est enzymes mapped unto EstX_Ec_ . Structural conservation, calculated and scored based on C-alpha RMSD values when comparing all four Est structures, is indicated by putty thickness (least conserved regions are shown by thicker putty radii). Sequence conservation, assessed via AL2CO scoring (40), coloured on spectrum of orange (least) to greencyan (most). **f**. EstX_Ec_ catalytic triad and oxyanion whole residues, coloured by sequence conservation. **g**. Complete macrolide esterase binding pocket mapped by substrate substituent localization: 5-pocket, ring-pocket, and 14-pocket.

Concomitant with the widespread use of macrolides has been a rise in resistance to these antibiotics (3, 6). Among the various mechanisms by which bacteria circumvent the effects of these antibiotics is enzyme mediated modification, which results in a loss of affinity for the 50S ribosome (3, 6). Until recently, two classes of macrolide resistance enzymes were known: erythromycin esterases and macrolide phosphotransferases (7, 8). The erythromycin esterases are a unique family of hydrolases that cleave the ester bond of macrolides with 14- and 15-membered rings. In contrast, macrolide phosphotransferases are evolutionarily related to protein kinases and aminoglycoside kinases and phosphorylate a hydroxyl group on one of the deoxy sugars that can be attached to 14-, 15-, or 16-membered lactones (7, 8).

We recently identified a third class of macrolide resistance enzymes, expanding the diversity of known resistance mechanisms for these antibiotics. Through functional annotation of a plasmid-encoded enzyme of unknown function carried by a multidrug resistant bacterium, *Sphingobacterium faecium*, from feedlot cattle water bowls, we discovered an antibiotic esterase, now named EstT_Sf_ (9). A retrospective analysis of genome and metagenome sequences also revealed that the *estT* gene is among the most abundant and frequently detected antibiotic resistance genes in cattle-associated bacteria (10). Consistent with the widespread use of macrolides, EstT homologs are present in many pathogenic and non-pathogenic bacteria across diverse biological niche (11-13). EstT_Sf_ and its homologs belongs to the α/β-hydrolase superfamily and uses a conserved Ser-His-Asp catalytic triad to hydrolyze the lactone ester bond. This catalytic mechanism is distinct from that of erythromycin esterases and employs a ping-pong bi-bi reaction involving an acyl-serine enzyme intermediate (14). Furthermore, in contrast to erythromycin esterases, the substrates for EstT enzymes are believed to be 16-membered ring-containing macrolides, such as tylosin (TYL) and tylvalosin (TYV) that are frequently used as veterinary drugs. Discovery of EstT_Sf_ also enabled the annotation of a long-named but only recently studied enzyme EstX as a macrolide esterase with similar activity (15, 16).

Here, we present functional and structural studies for four different macrolide Est enzymes: the original EstT identified in *Sphingobacterium faecium* (EstT_Sf_), two EstT homologs found in *Sphingobacterium thalpophilum* (obtained from a clinical human would isolate) and *Bacillus cereus* (EstT_St_ and EstT_Bc_), and the long-named EstX from *Escherichia coli* (EstX_Ec_). These macrolide esterases represent a diverse collection of Est enzymes that only share 44-66% sequence identity (Table S1). Therefore, findings from these studies will provide insights for the expanding Est class of macrolide esterases. Crucially, our results provide detailed information on the interactions between diverse Est enzymes and various macrolide antibiotics. This information is valuable for the development of next-generation macrolide antibiotics that are less susceptible to the continuing global rise in antibiotic resistance.

## Results

### Screening for macrolide esterases

To initiate functional characterization of the macrolide esterases, recombinant enzymes were produced, purified and evaluated in vitro. The ability of EstT_Sf_, EstT_St_, EstT_Bc_, and EstX_Ec_ to hydrolyze representative macrolides containing 14-, 15- and 16-membered rings was assessed using HPLC/MS (Figure S1; Figure S2). None of the enzymes were able to hydrolyze erythromycin (ERY) (14-membered) or tulathromycin (TUL) (15-membered) whereas enzyme-dependent hydrolysis of 16-memebered antibiotics was observed (Table 1). At 1 μM, all four enzymes hydrolyzed 200 μM of TYL and TYV within 4 h. In contrast, only EstT_Sf_ was able to completely hydrolyse tildipirosin (TIP) within 4 h, and the reaction of EstX_Ec_ with tilmicosin (TIL) also appeared to be slower than those catalyzed by the two other homologues. EstT_St_ and EstT_Bc_ possessed the most promiscuous activities in that hydrolysis of two leucomycins macrolides, spiramycin (SPM) and josamycin (JOS), was also observed. Collectively, the results are consistent with previous reports (9, 11, 13, 15). However, how the relative activities are reflected in bacterial phenotypes have not been compared. Accordingly, we assessed the capacity of these genes to protect *E. coli* from a series of macrolides, including tylosin-based antibiotics and leucomycins (Table S2). In lysogeny broth (LB) supplemented with 1 mM EDTA to increase membrane permeability (17), all 4 genes conferred resistance to TYL whereas only *estT*_Sf_ provided robust protection against semi-synthetic derivatives thereof. Additionally, modest protection by *estT*_St_ from a human clinical isolate was noted for SPM. Overall, the phenotypic results were consistent with qualitative measures of enzymatic activities and tentative assignment of macrolides as substrates or not.

**Table 1.** Qualitative analysis of macrolide esterase activities.

|  | EstT<sub>Sf</sub> | EstT<sub>St</sub> | EstT<sub>Bc</sub> | EstX<sub>Ec</sub> |
| --- | --- | --- | --- | --- |
| *14- 15-membered macrolides* |  |  |  |  |
| Erythromycin (ERY) | NS | NS | NS | NS |
| Tulathromycin (TUL) | NS | NS | NS | NS |
| *Leucomycin series* |  |  |  |  |
| Spiramycin (SPM) | NS | S | PS | NS |
| Josamycin (JOS) | NS | PS | PS | NS |
| *Tylosin series* |  |  |  |  |
| Tylosin (TYL) | S | S | S | S |
| Tylvalosin (TYV) | S | S | S | S |
| Tilmicosin (TIL) | S | PS | S | PS |
| Tildipirosin (TIP) | S | PS | PS | PS |

S: substrate (complete hydrolysis in 0.5-4 h); PS: poor substrate (complete hydrolysis after 4-8 h); NS: non-substrate (no hydrolysis after 24 h)

### Overall structure for macrolide esterases

Structural studies were pursued for all four macrolide esterases, with the intent to gain information on substrate recognition. For this we used variant enzymes which had the catalytic serine mutated to an alanine (S_cat_→A). Furthermore, all crystallization experiments contained either TYL or TYV. Five crystal structures of macrolide esterases EstT_Sf_, EstT_St_, EstT_Bc_ and EstX_Ec_ (2x) were solved at a resolution of 3.19 Å, 2.18 Å, 1.70 Å, 1.65 Å and 1.99 Å, respectively (Table S3). As anticipated, the three-dimensional structures confirmed the enzymes to belong to the α/β-hydrolase superfamily. Each Est enzyme adopts the canonical α/β-core fold comprised of an eight-membered β-sheet core and flanked by several α- helices on either side (Figure 1). Additional to the hydrolase α/β-core, is an immobile modular domain (18). Each Est-type crystal structure possesses a single cap domain, constructed of four α-helices and housed near the C-terminal end of the core β-sheet with large flexible loop regions linking the cap to the core domain (18). Towards the interface between the cap and core domains, the cap partially covers a pocket that extends into a large “canyon”-like fold which carves its way along the core domain’s surface (Figure 1g). Localized towards to the center, the conserved Ser-His-Asp catalytic triad can be found at positions: S126-H286-D258 (EstT_Sf_), S102-H260-D232 (EstT_St_), S102-H263-D235 (EstT_Bc_) and S102-H261-D233 (EstX_Ec_). Adjacent to the catalytic triad is the archetypical hydrolase oxyanion hole, conserved at positions A32 and L103 in all Est enzymes, but for EstT_Sf_ (A56, L127) (Figure 1f).

While the four Est structures share significant similarity, differences can be noted in specific regions (Figure 1e). For example, an ordered loop (L_11_) present in EstT_Sf_, EstT_St_, and EstX_Ec_ is presumably disordered in EstT_Bc_, as residues G212 – H221 could not be resolved in electron density maps. Similarly, an α-helix in present in EstT_Sf_, EstT_Bc_, and EstX_Ec_ was not observed in EstT_St_ given the lack of interpretable density for residues P138 – F152 in that enzyme. Interestingly, these two regions are located at the active site opining and could influence macrolide binding.

### Macrolide interactions with Est active sites

Even though, all crystallization experiments included either TYL or TYV, not all determined structures contained convincing electron density to place (part of) a macrolide in the active site. Polder maps for both EstT_Bc_ and EstX_Ec_ structures did allow for identifying TYL (EstX_Ec_) and TYV (EstT_Bc_ and EstX_Ec_) in the active sites (Figure 2). Noteworthy, electron density adjacent to the mutated Ala102 residues revealed that despite employing inactive variants for the structural studies, hydrolysis had occurred. This was not unexpected, as the timescale for crystallization was 3-4 weeks and mutations of the serine to an alanine in EstT_Sf_ (9) and other hydrolases that employ the catalytic triad have resulted in variants that still retain residual activity (19, 20) .Therefore, the complex structures represent the product bound states, with linearized TYL and TYV (L-TYL and L-TYV) found in the active sites.

**Figure 2.**
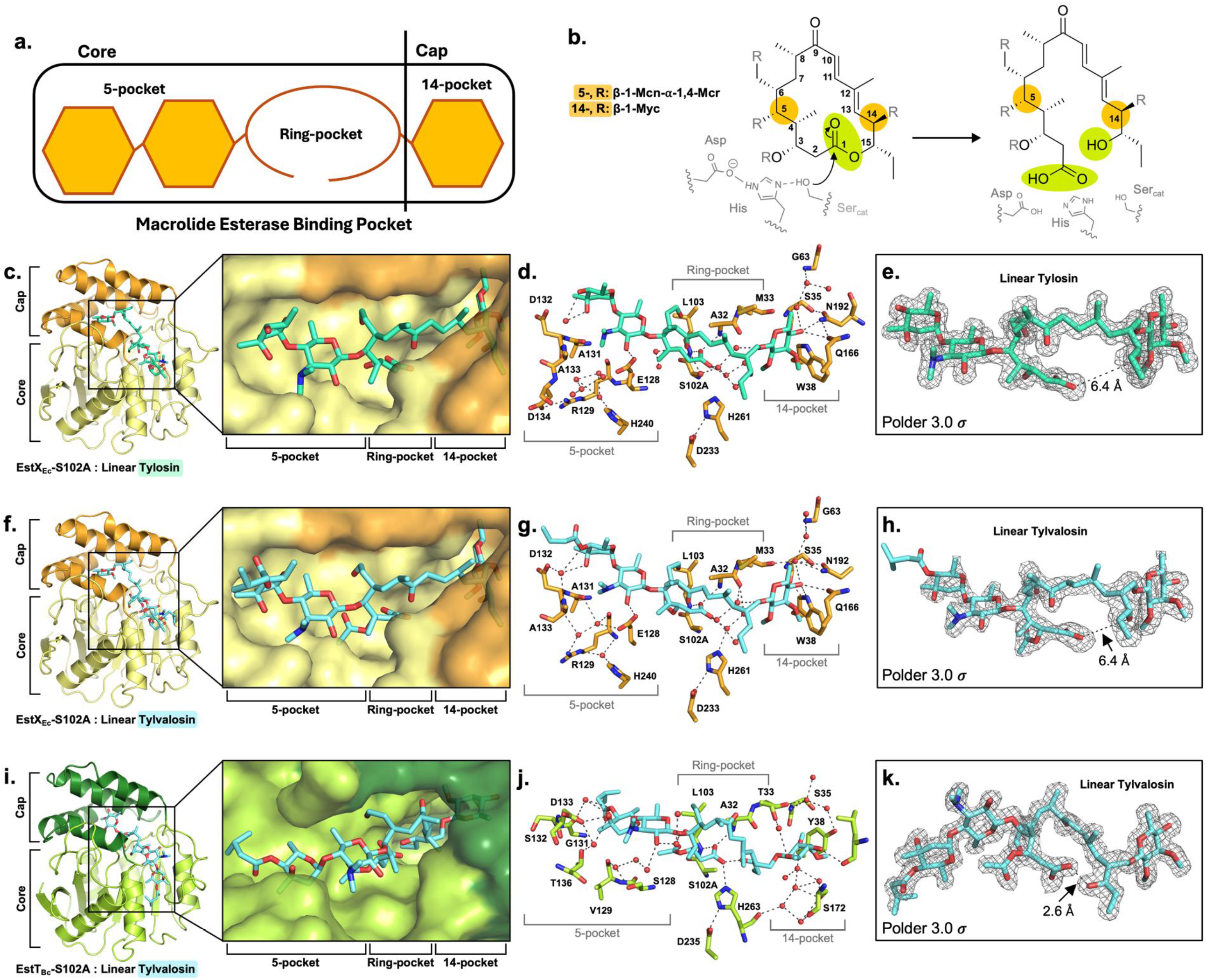
Ligand biding modes for EstX_Ec_ and EstT_Bc_. **a.** schematic representation of the macrolide esterase binding pocket regions, mapped according to 16-membered tylosin substitution nomenclature. **b**. Ser-His-Asp catalysis of 16-membered macrolide hydrolysis schematic. Substrate binding with EstX_Ec_ (yellow) is shown in complex with linear **c**. tylosin (greencyan) and **f**. tylvalosin (aquamarine). **i**. EstT_Bc_ (green) is shown in complex with linear tylvalosin. **d**., **g**., **j**. specific 5-, ring-, and 14-pocket residues involved in water-cage binding of substrate. Polder map densities describe linear **e**. tylosin and **h**. tylvalosin bound to EstX_Ec_ and **k**. tylvalosin bound to EstT_Bc_.

The three Est complex structure all exhibit a similar binding mode for a linearized macrolide. The two deoxysugar moieties that are attached to the 16-membered cleaved rings at positions 5 and 14 in both L-TYL and L-TYV are located on either end of the large and narrow α/β-hydrolase active site. In EstT_Bc_ and EstX_Ec_ bound structures, the linearized macrolides are oriented such that the 5-position disaccharide (β-1-mycaminose-α -1,4-mycarose) (21) faces the active sites’ mouth along the hydrolase core. The 14-position monosaccharide substituent (β-1-mycinose group) (21) sits within the enclosed pocket beneath the cap domain. Thus, the substrate binding site can be viewed as consisting of three sub-sites, the 5-pocket, the lactone ring-pocket, and the 14-pocket. (Figure 1g, 2a).

In the 5-pocket, the binding poses for di-deoxy sugar moiety reveal some variability, while the mycaminose (Mcn) sugar groups in L-TYL and L-TYV at position 5 consistently binds in the same orientation, the second terminal mycarose (Mcr) group, also present in both linearized macrolides, has two distinct poses. The two different poses appear not to be linked to the difference between TYL and TYV, i.e. TYV has an additional acetyl-4’-isovaleryl group on the 3-postion of the Mcr moiety but are likely caused by structural differences between EstT_Bc_ and EstT_Ec_. As mentioned above, the L_11_ loop is disordered in EstT_Bc_, permitting a differing orientation of the Mcr sugar. In the ring-pocket, the linearized former 16-membered lactones are positioned such that carboxylic acid groups created during hydrolysis are in close proximity to the catalytic triad residues and the oxyanion amides. The hydroxyl arms appear to be rotated outwards and away from the carboxyl-(i.e., where the substrate bound state’s ester bond is presumed to bind). The distance between the carboxylic acid group and hydroxyl group are 2.6 Å in EstT_Bc_ and 6.4 Å in the two EstX_Ec_ complex structures (Figure 2e,h,k). Noteworthy, that the carboxylic acid groups appear to be stationary while the hydroxyl arms move away, is consistent with the canonical esterase reaction mechanism for α/β-hydrolase. Finally, the 14-pockets are slightly different between EstT_Bc_ and EstX_Ec_ such that the poses for the mycinose (Myc) moieties are different between these two esterases, with the difference being a rotation of the sugars.

It should be noted that the nature of the interactions observed between the linearized macrolides and the Est enzymes observed in the three product state structures is highly unusual. Surprisingly, there are very few hydrogen bonds between L-TYL and L-TYV with the two Est enzymes (Table S4). Given that each linearized macrolide has the potential to make over 20 specific hydrogen bond interactions, it is unexpected that the Est enzymes only have 5-6 hydrogen bonds with the product (Table S4). However, compensating for this are additional water mediated hydrogen bonds that stabilize the observed product state complexes (Figure 2d,g,j). It is tempting to speculate that these esterases employ a water-cage to capture a more diverse collection of substrates then if direct specific hydrogen bonds were used by these enzymes. An additional observation is that in the ring-pocket, the predominant interactions are hydrophobic in nature.

### Steady-state kinetics corroborate expected AMR phenotype

Structural insights and varied AMR phenotypes prompted further characterization of the two most distinct homologues in the set: EstT_Sf_ and EstX_Ec_. Steady-state kinetic parameters for the reactions of EstT_Sf_ and EstX_Ec_ with tylosin, the presumed cognate substrate of each, were measured by single-injection ITC methods (Table 2; Figure 3; Figure S3). The reaction of EstT_Sf_ with TYL was 5-fold more specific than that of EstX_Ec_. Moreover, EstX_Ec_ was slow to turnover to TIL, which was categorized as a poor substrate during screening. In this case, insufficient heat of reaction between EstX_Ec_ and TIL required the use of HPLC/MS to determine an apparent *k*_cat_ ∼11 h^-1^ for the reaction. This slow turnover was commensurate with the qualitative observation of more TIL than linear product after 8 h of reaction (Figure S4). In contrast, the apparent turnover of TIL by EstT_Sf_ was on the same timescale as TYL, corroborating both the qualitative biochemical analysis and protection conferred when expressed in *E. coli*. Interestingly, turnover was not limited by binding as dissociation constants for the EstTs, EstX_Ec_, variants were between 10 to 70 μM for TYL and TIL (Table 3, Table S5). More broadly, similar equilibrium binding observations could be made between enzymes and macrolides that were or were not substrates (e.g., EstT_Sf_:TIP and EstT_Bc_:TIP), suggesting that the observed water-cage indeed enables promiscuous binding.

**Table 2.**
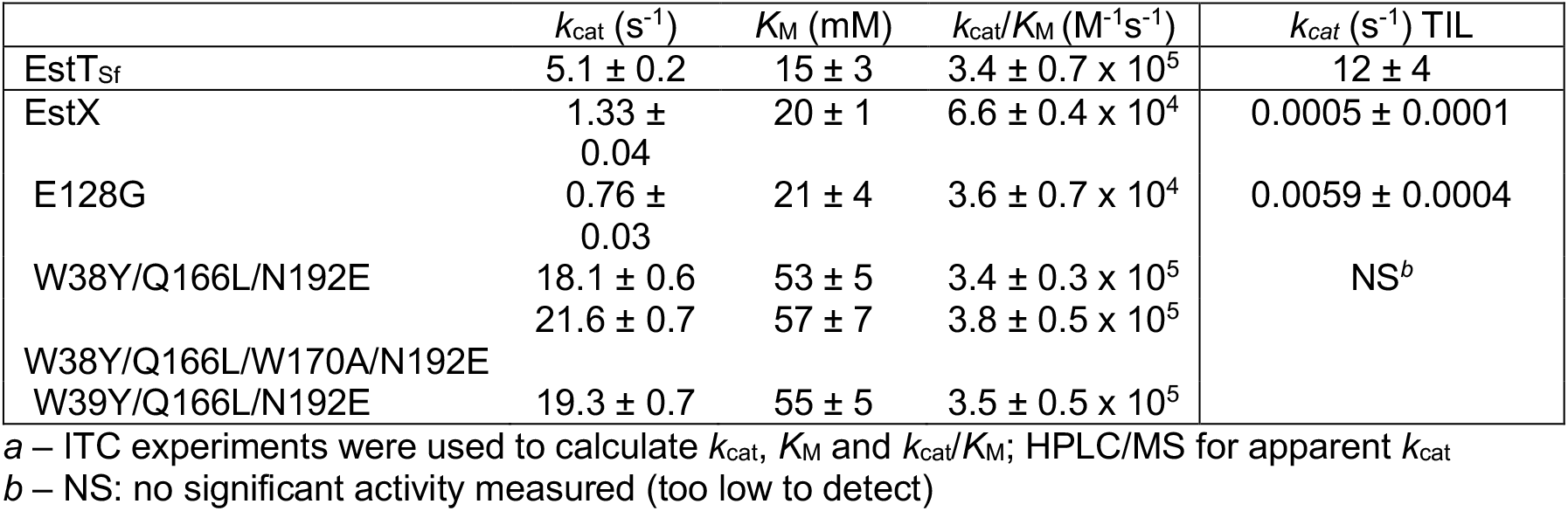
Steady-state kinetic parameters determined for the reaction of EstT, EstX and variants with TYL and TIL^*a*^.

**Table 3.** Dissociation constants (*K*_D_, mM) measured for catalytically inactive esterases and macrolides.

| S<sub>cat</sub>→A variant | TYL | TYV | TIL | TIP |
| --- | --- | --- | --- | --- |
| EstT<sub>Sf</sub> | 21 ± 5 | 21 ± 3 | 25 ± 3 | 62 ± 13 |
| EstT<sub>St</sub> | 22 ± 3 | 21 ± 5 | 26 ± 3 | 47 ± 8 |
| EstT<sub>Bc</sub> | 57 ± 7 | 90 ± 40 | 45 ± 6 | 39 ± 4 |
| EstX<sub>Ec</sub> | 10 ± 2 | 125 ± 9 | 55 ± 14 | 86 ± 15 |

**Figure 3.**
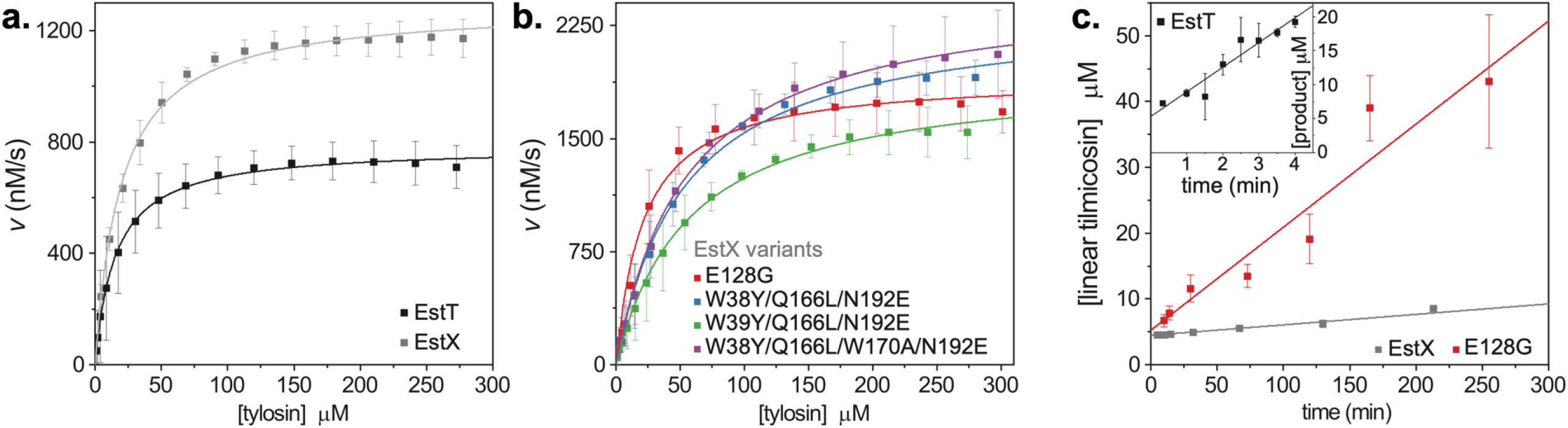
Steady-state kinetic analysis of EstT_Sf_ and EstX_Ec_. Michaelis-Menten plots showing **a.** turnover of tylosin by EstT_Sf_ and EstX_Ec_ and **b**. EstX_Ec_ variants inspired by EstTSf. **c**. Linear plots of time-dependent tilmicosin hydrolysis by EstT_Sf_ (inset), EstX_Ec_ and EstX_Ec_ E128G.

We next used a structural comparison to incorporate site specific EstT_Sf_ character into the active site of EstX_Ec_. Four EstX_Ec_ variants – E128G, W38Y/Q166L/N192E, W39Y/Q166L/N192E and W38Y/Q166L/W170A/N192E – were generated based on observed proximity and interactions with the mycaminose and mycinose ring substituents in product-bound structures. The E128G variant and combinatorial variants are in the 5- and 14-pocket, respectively. The enzymes were active (Figure S3) and substitutions to EstX_Ec_ at the 14-pocket increased the specificity of the reaction with TYL (Table 2). In fact, the apparent second order rate constants for these reactions matched that observed between EstT_Sf_ and TYL for the 14-pocket variants. In contrast, the EstX_Ec_ E128G variant was relatively unchanged with respect to TYL hydrolysis, however, it displayed a 10-fold increase in turnover of TIL. Nevertheless, none of the variants displayed increased phenotypic resistance to TIL (Table 2) and TIL hydrolysis by the 14-pocket variant W38Y/Q166L/N192E was abrogated (Figure S4). Kinetic characterization of these variants suggests that the catalytic determinants of individual Est class of macrolide esterases are not simply interchangeable.

### *In silico* molecular modeling of macrolide substrate binding

In order to help rationalize the functional studies on Est macrolide interactions based on the structural data, molecular models were constructed for each of the four Est enzymes in complex with both substrates and non-substrate macrolides. All macrolides from both the tylosin and leucomycin series could be modeled into all four Est enzymes. This is somewhat curious as we show that some of the leucomycin antibiotics are not hydrolyzed by some of the enzymes. However, it does agree with the measured dissociation constants. Again, as alluded to above, it suggests that binding is not the only prerequisite for catalysis. The 14-membered ERY and 15-membered TUL macrolide antibiotics could not be modeled into the active sites of any of the Est enzymes. The principal reason was that these antibiotics have deoxy sugar extensions at the 3-position of the macrolactone which, when modeled into the active sites make severe steric clashes, thus providing a convincing rational for why these antibiotics are not hydrolyzed.

## Discussion

Since its discovery in 2023, macrolide esterases that belong to the α/β-hydrolase family have now been found in numerous bacteria, some of which are environmental, many are veterinary pathogens, and some are also human pathogens (9, 11, 13, 15). The studies presented here for a diverse set of four Est enzymes, that include both veterinary and human pathogens, demonstrate that these enzymes can hydrolyze 16-membered macrolide antibiotics of the tylosin series and confer resistance in a model system. Moreover, two of the four enzymes have an expanded substrate spectrum and can also confer resistance to macrolides belonging to the leucomycin series. The catalytic efficiencies of the EstT_Sf_ and EstT_Ec_ with their cognate substrate are one to two orders of magnitude greater than those reported for the disparate erythromycin esterase family (22). As both tylosin and leucomycin series include antibiotics that are currently used to treat bacterial infections in veterinary and human medicine, Est mediated antibiotic resistance is thus a growing One Health issue (23).

### Est macrolides esterases are a distinct class of α/β-hydrolases

While the realization that Est mediated macrolide resistance is a One Health problem is recent, the enzymes themselves have been present in the environment long before the use of macrolide antibiotics as is evident from the sequence diversity among the enzymes and diversity of niches in which they are found. There are seemingly countless EstT homologues with divergent sequences that are well-dispersed across the Bacteria (9, 11-13). In contrast, those named EstX are characterized by limited sequence diversity and an exclusive presence in few γ-Proteobacteria, including *E. coli, Klebsiella pneumoniae and Salmonella enterica* (15). Interestingly, despite the significant variation in amino acid sequences, as indicated by the % identity of ∼50%, the three-dimensional structures we have determined are remarkably similar with r.m.s.d. values only around ∼1 Å (Table S1).

Searching for structurally similar enzymes, using the DALI server, reveals that even though the α/β-hydrolase family, is extensive, with more than 3000 experimentally determined structures, the macrolide esterases are distinct from all of these (24, 25). The nearest structural homologues include other bacterial enzymes which perform a variety of common α/β-hydrolase reactions, such as: aclainomycin methylesterase RdmC (PDB: 1Q0Z), lipase ORF17 (PDB: 8PZG), esterase Ade1 (PDB: 8V16), epoxide hydrolase IbcK (PDB: 9C2G), and C-C hydrolase DxnB2 (PDB: 4LXG) (Table S6, Figure S5) (26-30). However, comparison shows that they have crucial differences with the Est enzymes presented here. Specifically, there is much variability in the cap domain, which is a major determinant for dictating substrate specificity in this family. The nearest structural homologs of the macrolide esterases all have cap domains elements in place of the Est-type 5-pocket region which would prevent the large macrolide substrates from binding to those enzymes (Figure S5). Indicating a uniqueness to the α/β-hydrolase macrolide esterase binding pocket. This, together with the limited sequence conservations, suggests that macrolide esterases are an ancient distinct branch of the α/β-hydrolase family tree.

### Macrolide specificity is not only dictated by binding

The observed macrolide specificity for the four esterases is surprising. For example, for EstT_Sf_ the results showing SPM is a good substrate while JOS is a poor substrate is difficult to rationalize based on the minor chemical differences between the two macrolides. The addition of the acetyl-4’-isovaleryl group on the 3-postion of the Mcr moiety in JOS should be readily tolerated by EstT_Sf_ based on the ability of this enzyme to cleave both TYL and TYV. Also, the absence of a deoxy sugar on the 9-position in JOS vs. SPM, should not be detrimental as all other substrates for this enzyme lack that moiety. In fact, based on chemical similarity one would predict that JOS is a better substrate for EstT_Sf_ then SPM, but the opposite is observed. Intriguingly, the difficulty in rationalizing macrolide specificity is exacerbated by the *in-silico* modeling studies. While the amino acid conservation in the active site for the four macrolide esterases is limited (Figure 1), all six 16-membered macrolides studied could be placed in the active sites of all four enzymes without steric clashes between the enzymes and the macrolides. The principal reason that modeling of the complex states could be readily accomplished is that, as shown in the experimental structures with L-TYL and L-TYV, the macrolides are largely bound to the enzymes through a malleable water-cage for the 5- and 14-position extensions and through non-specific hydrophobic interactions for the macrolactone ring. These modeling results suggest that the observed macrolide specificity is unlikely to be the result of the enzymes’ inability to bind macrolides, leading us to conclude other factors must be impacting catalysis. The idea that binding is perhaps not the crucial factor in dictating catalysis is supported by the dissociation data presented (Table 3). These results show that whether a macrolide is a good or poor substrate is not dependent on the affinity, as these values remain remarkably constant regardless of the ability of the enzyme to cleave the macrolide.

An alternative rationalization for the differences in macrolide specificity could be that macrolides may bind to the enzymes, but that the binding mode is not always conducive to catalysis. It is well established that the exact conformation of the substrate bound state is crucial for catalysis and that minor shifts in the placement of substrate atoms in relation to catalytic residues have dramatic impacts on reaction rates (31). The limited number of specific interactions that we have observed between macrolides and Est enzymes, specifically in the ring-pocket, suggests that exact placement of the ester bond is expected to be sufficiently variable between the different Est-macrolide substrate complexes, precipitating differing reaction rates. Unfortunately, the precise position of the ester bond would likely be dictated by subtle differences in the enzymes the substrates and the fluid nature of the water-cage that would be difficult to predict given the current accuracy of computational modeling methods. The difficulty to rationally predict which factors influence macrolide specificity is demonstrated by our efforts to modify EstX_Ec_ to accept TIL as a *bona fide* substrate by mutating key residues to those observed in EstT_Sf_ (Table 2). While we were able to alter catalytic properties, we were unable to increase the efficiency of TIL cleavage. This implies that we currently are unable to propose a predictive model for 16-membered macrolide substrate specificity for the macrolide esterases studied here. However, our results suggest that macrolide esterases are promiscuous in that they can bind tylosin and leucomycin macrolides, but that binding is not always followed by hydrolysis, as this requires optimal placement in the active site, which is not always achieved due to the lack of specific interactions. Thus, in a perverse way, the water-cage structure is both responsible for expanding and limiting the macrolide specificity for these α/β-hydrolase by simultaneously allowing for promiscuous binding and preventing optimal binding.

### Implications for addressing macrolide antibiotic resistance

Our results suggests that macrolide esterases belonging to the α/β-hydrolase family are promiscuous binders of 16-membered macrolide antibiotics, and that members of this class can confer resistance to natural and semisynthetic drugs. Given the expanding diversity of Est enzymes, it is near inevitable that Est variants can be found with greater substrate spectrum and enhanced catalytic activities. This raises profound concerns for the continued use of these antibiotics in both veterinary and human settings. Moreover, the ability of these enzymes to already provide resistance to semisynthetic macrolide variants, indicates that pursuing the development of next generation macrolides might prove challenging. Specifically, given the promiscuous binding properties we believe Est esterases possess, significant changes to the 16-membered lactone scaffold might be required to thwart the resistance properties of these enzymes.

However, the structures for the product state and models for the substrate-bound states provide insights into which avenues in macrolide drug development are more promising. The similarity in binding poses for the six different macrolides to the four diverse esterases suggests that alterations to the core 16-membered lactone scaffold that interferes with binding to one of the enzymes will very likely also interfere with binding to all macrolide esterases. For example, possible expansions on the scaffold by adding bulky groups at position 3 or 12 would likely result in a macrolide variant that will be unable to bind to any Est enzymes. Obviously, any changes to the chemical structure of the macrolide should not interfere with its ability to bind to the ribosome so as not to impact its antibiotic properties. In our example this would actually disqualify any extensions at the 12-position, based on the structure of the bacterial ribosome with tylosin (Figure 4) (31). Nonetheless, adding extensions at the 3-position might be a potential avenue. This is supported by our results for ERY which is able to bind to the ribosome, but which is not a substrate for macrolide esterases, in part because of its deoxy sugar at the 3-position. Of course, other avenues for next generation macrolide development can be envisioned guided by the results presented.

**Figure 4.**
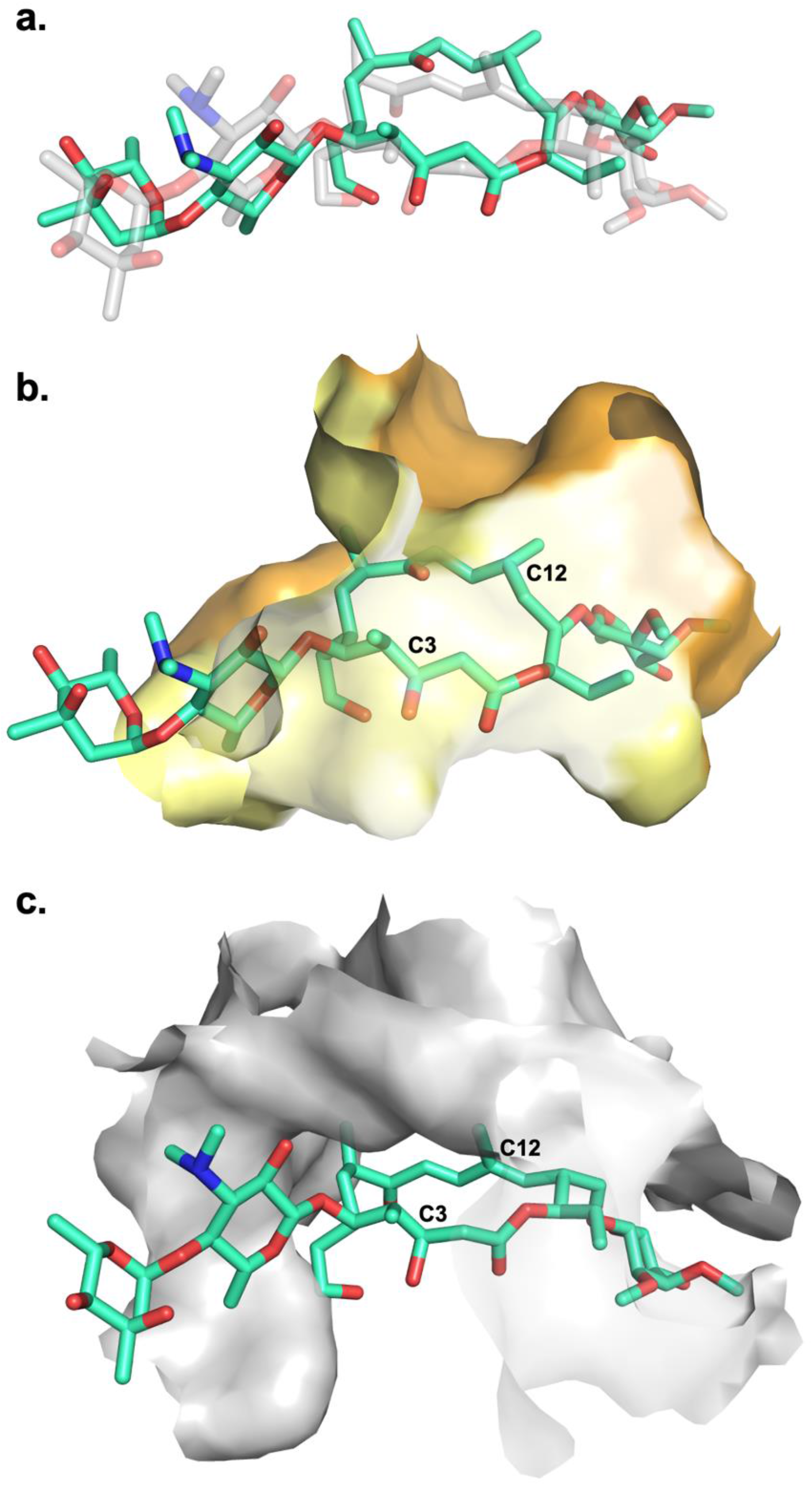
Comparison between cyclic tylosin binding to EstX_Ec_ and the ribosome. **a.** Superposition of modeled tylosin (greencyan), in the conformation modeled with EstX_Ec_ (yellow), and tylosin (transparent grey) in the conformation observed in the 50S ribosomal subunit (PDB: 1K9M) (26). Conserved tylosin binding modes are compared inside the **b**. Est-type (EstX_Ec_) and **c**. ribosome 50S (grey) binding pockets.

## Materials and Methods

### Reagents

All reagents were of commercial origin. Vendors and catalogue numbers for specific antibiotics are listed: erythromycin (MilliporeSigma, E5359,) tulathromycin (MedChemExpress, HY-15662), spiramycin (Sigma-Aldrich, S9132), josamycin (MedChemExpress, HY-B1920), tylosin tartrate (ThermoScientific, AC463070050), tilmicosin (MedChemExpress, HY-B0905), tildipirosin (LGC Standards, 77678-842), tylvalosin tartrate (Cayman Chemical, 33849).

### Protein production

EstT_Sf_(UXD71803.1), EstT_St_ (WP_028071475.1), EstT_Bc_ (MFC8766929.1), EstX_Ec_(PP210101.1) and variants were cloned in pET29b(+), produced and purified as previously described with minor changes (9). To simplify production of EstT_Sf_, the first 24 codons were omitted from the construct. The proteins were produced in *E. coli* Rosetta (DE3) or BL21(DE3). Cultures were grown to an optical density at 600 nm of 0.6 before recombinant gene expression was induced with isopropyl β-D-1-thiogalactopyranoside (IPTG), followed by overnight incubation at 16-25 °C with shaking at 200 rpm. Cells were harvested by centrifugation at 6000 g, resuspended in 20 mM TRIS, 300 mM NaCl, pH 7.5, and lysed at 25,000 PSI. Lysates were clarified by centrifugation at 31,000 g for 20 min at 4 °C and filtered through a 0.45 µm membrane before loading onto a gravity column packed with Ni-NTA resin. Non-tagged proteins were washed stepwise from the column with increasing concentration of imidazole, and macrolide esterase-containing fractions were pooled, concentrated, and buffer-exchanged into 100 mM potassium phosphate buffer (KPi), pH 7.0, using 10 kDa MWCO centrifugal filters. Protein aliquots were flash frozen and stored at -80 °C until use.

For structural biological studies this protocol was modified to address the increased quantity and purity needed for those experiments. Est proteins were expressed in BL21 (DE3) *E. coli* in 1L culture flasks containing autoinducing ZYP-5052 expression media at 37°C for 3 hr before overnight expression 16°C (16 hr expression time). Cells were lysed by sonification (Branson 450 Digital Sonifier, amplitude 60% for 10 min), and non-soluble debris spun down for 20 min at 18,000 g. Protein purification was further adapted by adding the following additional steps, after Ni-NTA column. Fractions containing Est protein were pooled and treated with Thrombin digestion (via Thrombin cleavage column). Resultant protein fractions were pooled and concentrated in 15 mL Amicon® Ultra Centrifugal Filters (10 kDa cut-off), buffer exchanged (50 mM Bis-Tris, pH 6.7), and passed through a 0.22 mm filter before purification via anionic exchange (MonoQ HR 16/10), eluting on a salt gradient (final buffer concentration: 500 mM NaCl). Following anionic exchange, Est proteins were purified with a final gel filtration step (Superdex75 26/60; 20 mM HEPES, pH 8, 150 mM NaCl for EstT_Bc_ and 20 mM Tris, pH 8.0, 100 mM NaCl for all others), and stored at -20°C.

### Qualitative biochemical assessment of macrolide esterases

Macrolide esterases (1 μM) and putative substrates (200 μM ERY, TUL, SPM, JOS, TYL, TYV, TIL and TIP) were reacted in 100 mM KPi, pH 7, at room temperature (RT) for defined time intervals (minutes to hours) (Figure S3). Reactions were quenched with 1% acetic acid (v/v), dried in vacuo, resuspended in acetonitrile and filtered prior to HPLC/MS analysis. Chromatographic separation of hydrolytic products and substrates was performed using a Phenomenex Kinetex F5 column (100 Å, 5 μm, 100 × 4.6 mm) with a binary solvent system as previously described (9).

### MIC determinations

To assess protection conferred by putative macrolide esterases, synthetic, *E. coli* codon optimized *estT*_*Sf*_, *estT*_*St*_, *estT*_*Bc*_ and *estX*_*Ec*_ genes and variants, were placed downstream of a proD-insulated promoter and the Bba-B0032-RBS (32). The *his* operon terminator followed the gene, which was cloned into a pTWIST Amp^R^ high-copy vector (Twist Bioscience). An *ereA* gene-carryingc construct (protein ID: AAO38247) was prepared as a second control with a known AMR phenotype. *E. coli* DH5α transformants were propagated in LB with or without 1 mM EDTA to log-phase before dilution to an optical density of 5 x 10^-5^ in 200 μL aliquots of the same media containing 2-fold dilution series of macrolides in 96-well microtitre plates. DMSO was used to prepared macrolide stocks and included at 1-2.5% (v/v) in the assays. Bacterial growth was assessed after overnight incubation at 37 °C without shaking, measuring by absorbance at 600 nm using a BioTek Synergy HTX Multimode Reader (Agilent). The MIC (Table S7, Table S8) was defined as the lowest macrolide concentration that prevented growth in three independent biological measurements.

### Crystallization and X-ray diffraction data collection

To enable crystallographic structure determination of Est enzymes in complex with macrolides, all enzymes used for these studies were inactive through mutation of the catalytic serine to an alanine (S_cat_→A). Est crystals were grown in INTELLI-PLATE® 96-3 Low Volume Reservoir plates, at 4°C, by sitting-drop vapour diffusion. 0.6 µL drops contained a 1.1 mixture of mother liquor and 5-10 mg/ml protein solution in storage buffer that also contained 4-10 mM of either TYL or TYV macrolide. The used mother liquor for the various Est crytals were: EstT_sf_ - 0.1 M HEPES pH 7.5, 10% PEG 4K; Est_St_ - 0.1 M Tris, pH 9.6, 0.2 M MgAc, 20% PEG 3350; EstT_Bc_ - 0.1 M MES, pH 6.5, 0.1 mM MgAC, 12% PSM, 10% ethylene glycol; EstX_Ec_ - 0.1 M sodium cacodylate, pH 5.3, 0.1 mM tacsimate, 10% PEG, 15% PSB (TYL crystallization), and 0.1 M Bicine, pH 9.5, 22% PSB (TYV crystallization). To enable data collection under cryogenic conditions, crystals were briefly soaked in a cryoprotectant solution composed of mother liquor and 25% PEG 4K or 20% glycerol and flash frozen in liquid nitrogen. Diffraction EstT_St_, and EstX_Ec_ datasets were collected at the Canadian Macromolecular Crystallography Facility Beamline 08B1-1 at the Canadian Light Source (Saskatoon, SK, Canada). Diffraction data for EstT_Bc_ and EstT_Sf_ were collected at the Advanced Photon Source 23-ID-B Beamline, APS (Lemont, IL, USA). The data were processed using the HKL2000 suite of programs (Table S3) (33).

### Structural determination and refinement

Crystal structures for the four macrolide Est enzymes were solved by molecular replacement using the PHENIX software package, and search models generated by AlphaFold (34, 35). Model refinement was carried out in successive rounds of real- and reciprocal-space refinement using PHENIX and COOT (36). Model modifications, corrections, and additions were based on 2F_o_-F_c_ and F_o_-F_c_ electron density maps. While all crystallization experiments contained either TYL or TYV and used inactive mutants, no convincing density for macrolides were found in EstT_Sf_ and EstT_St_, and hence these structures are designated as apo structures. Electron density that unambiguously belonged to macrolides were observed in the active sites of EstT_Bc_ and EstX_Ec_. However, despite that the inactive S_cat_→A mutations, polder maps convincingly showed that macrolide ester bonds were cleaved in these structures. As the product of the reaction are linearized macrolides, we refer to these products here as L-TYL and L-TYV. Model refinements were performed until no significant improvements could be achieved, judged by R/R_free_ while maintaining reasonable stereochemistry (Table S1).

### Enzyme kinetic studies

Steady-state kinetic parameters for the reaction of EstT_Sf_, EstX_Ec_ and variants with TYL were determined using single-injection ITC measurements. Continuous reaction monitoring was enabled by a Nano-ITC LV isothermal titration calorimeter (Malvern Panalytical). The sample cell was filled with degassed enzyme solutions in 100 mM KPi, pH 7.0, and reactions were initiated by the injection of TYL with the syringe stirring speed set to 200 rpm and thermal power was recorded at 2 s intervals. The final concentration of TYL in each reaction was 389 µM and the concentration of enzymes was optimized to ensure both saturation and that reactions were complete within 6 min. Control experiments included buffer injections into enzyme or TYL injections into buffer. Time-resolved thermogram data was transformed to generate Michaelis-Menten plots as previously described (37, 38, 39).

Reactions of esterases with 400 µM TIL were performed in 100 mM KPi, pH 7.0 at RT and quenched with 1% acetic acid (v/v) at defined time intervals (seconds to hours) to determine an apparent *k*_cat_ values. Product formation was monitored using a Phenomenex Kinetex F5 column (100 Å, 5 μm, 100 × 4.6 mm) and a two-solvent gradient of solution A (water, 0.05 % formic acid) and solution B (acetonitrile, 0.05 % formic acid). LC-ESI-MRM measurements were conducted with a Q-Sight 420 LC-MS/MS Triple Quadrupole instrument (PerkinElmer) (Figure S6, Table S9). Quantification was enabled by a standard curve of linear TIL.

### Equilibrium binding studies

Dissociation constant (*K*_d_) values (Table 3, Table S5, Figure S7) were determined for enzyme:macrolide pairs using catalytically inactive of EstT_Sf_, EstT_Bc_, EstX_Ec_ S_cat_→A and additional specific variants. Binding of macrolides was measured by monitoring changes to intrinsic tryptophan fluorescence (λ_ex_ = 280 nm, λ_em_ = 350 nm). Inactive enzymes (140 nM) were mixed with macrolide (7 to 450 μM) in 100 mM KPi, pH 7.0 in 200 μL volumes within microtitre plates. DMSO was used to facilitate solubilization of macrolides.

### Molecular modeling of macrolide binding

*In silico* molecular modeling of macrolides in the substrate bound state was performed for all four macrolide Est enzymes with the following antibiotics: SPM, JOS, TYL, TYV, TIL, and TIP. First, models were generated for EstT_Bc_ with TYL and EstX_Ec_ with TYL and TYV, based on the crystal structures of these enzymes with L-TYL and L-TYV (Figure 4a). Ring closure could readily be achieved without altering most of the macrolide structure and while enforcing correct stereochemistry. The ester bond could be placed in the correct position for catalysis, requiring only atoms downstream of the deoxy sugar moiety at the 14-positions to be repositioned. Based on these models the remaining macrolides were modeled into the EstT_Bc_ and EstX_Ec_ active sites by adding and/or removing substituents on the TYL and TYV models. Macrolide bound states for EstT_Sf_ and EstT_St_ were subsequently generated using those for EstT_Bc_ and EstX_Ec_ as blueprints. We also attempted generating substrate bound states for the non-substrates ERY and TUL, to further aid in rationalizing substrate specificity.

## Data Availability

Coordinates and structure factors of EstT_Sf_, EstT_St_, EstT_Bc_, EstX_Ec_:L-TYL, and EstX_Ec_:L-TYV have been deposited in the Protein Data Bank with accession codes: 11TS, 11TT, 11TU, 11TV, and 11TW, respectively. EstT_Sf_ (UXD71803.1) was used as a blastp query (nr protein sequence database) to access EstT_St_ (WP_028071475.1) EstT_Bc_ (MFC8766929.1) and EstX_Ec_ (PP210101.1). Additional data supporting the findings of this study are available from the corresponding authors upon reasonable request.

## Acknowledgments

We are grateful to the University of Saskatchewan Protein Characterization and Crystallization Facility for access to an ITC and the Saskatchewan Structural Sciences Centre for access to a mass spectrometer provided by Perkin Elmer. X-ray diffraction data collection was carried out at the Canadian Light Source (CLS) CMCF-BM beamline and the GM/CA Advanced Photon Source (APS) 23-ID-B beamline. CLS is a national research facility of the University of Saskatchewan which is supported by the Canada Foundation for Innovation, the Natural Sciences and Engineering Research Council, the National Research Council, the Canadian Institutes of Health Research, the Government of Saskatchewan, and the University of Saskatchewan. GM/CA@APS has been funded by the National Cancer Institute (ACB-12002) and the National Institute of General Medical Sciences (AGM-12006, P30GM138396). This research used resources of the Advanced Photon Source, a U.S. Department of Energy (DOE) Office of Science User Facility operated for the DOE Office of Science by Argonne National Laboratory under Contract No. DE-AC02-06CH11357. We also thank past and present members of both the Ruzzini and Berghuis Labs for their help and support, in particular: Jyllian Jackson of the Berghuis Lab, who participated in the research through the Indigenous Mentorship and Paid Research Experience for Summer Students (IMPRESS) program. This work was funded by Beef Cattle Research Council (ANH.01.22), Results Driven Agriculture Research Fund (2024F2477R), SK Agriculture Development Fund (20230048) and CIHR grant PJT-192008 to AR, and CIHR grant PJT-162365 to AMB. AMB is a member of the Centre de recherche en biologie structurale (CRBS), which receives funding from Fonds de Recherche du Québec (Health Sector) Research Centres Grant #288558. ETRK is a recipient of a FMHS Studentship in partnership with CAN-AMR-NET. IM is the recipient of an NSERC CGS-D.

## References

1. Hutchins, M.I., Truman, A.W. & Wilkinson, B. Antibiotics: past, present and future. Curr. Opin. Microbiol. 51, 72–80 (2019).

2. Hamad, B. The antibiotics market. Nature Reviews Drug Discovery 9, 675–676 (2010).

3. Golkar, T., Zieliński, M. & Berghuis, A.M. Look and outlook on enzyme-mediated macrolide resistance. Front. Microbiol. 9, 1942 (2018).

4. Dinos, G.P. The macrolide antibiotic renaissance. PJB 174, 2967–2983 (2017).

5. Poehlsgaard, J. & Douthwaite, S. The bacterial ribosome as a target for antibiotics. Nat. Rev. Microbiol. 3, 870–881 (2005).

6. Leclercq, R. Mechanisms of Resistance to Macrolides and Lincosamides: Nature of the Resistance Elements and Their Clinical Implications. Clin. Infect Dis. 34, 482–492 (2002).

7. Zieliński, M., Park, J., Sleno, B. & Berghuis, A. M. Structural and functional insights into esterase-mediated macrolide resistance. Nat. Commun. 12, 1732 (2021).\

8. Fong, D. H., Burk, D. L., Blanchet, L., Yan, A.Y. & Berghuis, A. M. Structural basis for kinase-mediated macrolide antibiotic resistance. Structure 25, 750-761.e5 (2017).

9. Dhindwal, P., et al. A neglected and emerging antimicrobial resistance gene encodes for a serine-dependent macrolide esterase. PNAS 120, e2219827120 (2023).

10. Kos, D., Jelinski, M. & Ruzzini, A. Retrospective analysis of antimicrobial resistance associated with bovine respiratory disease. Appl. Environ. Microbiol. 91, e0190924 (2025).

11. Xin, A., et al. An omics-based framework for investigating the emerging antibiotic resistance gene: The case of estT. Microbiol. Res. 303, 128377 (2026).

12. Zhou, Y., et al. Global distribution of alpha/beta hydrolase family macrolide esterases in Gram-positive bacteria. ISME J. 19 (2025).

13. Toa, H., et al. Functional characterization of macrolide esterase from cyanobacteria and their potential dissemination risk. NPJ Antimicrob. Resist. 4, 10 (2026).

14. Rauwerdink, A. & Kazlauskas, R. J. How the Same Core Catalytic Machinery Catalyzes 17 Different Reactions: the Serine-Histidine-Aspartate Catalytic Triad of alpha/beta-Hydrolase Fold Enzymes. ACS Catal. 5, 6153–6176 (2015).

15. Lin, J., et al. The global distribution of the macrolide esterase EstX from the alpha/beta hydrolase superfamily. Commun. Biol. 7, 781 (2024).

16. Dhindwal, P., Myziuk, I. & Ruzzini, A. Macrolide esterases: current threats and opportunities. Trends Microbiol. 31, 1199–1201 (2023).

17. Leive, L. A Nonspecific Increase in Permeability in Escherichia Coli Produced by Edta. Proc. Natl. Acad. Sci. U S A. 53, 745–750 (1965).

18. Bauer, T. L., Bechholz, P. C. F. & Pleiss, J. The modular structure of a/b-hydrolases. FEBS J. 287, 1035–1053 (2020). doi:10.1111/febs.15071.

19. Krishnan, R., Sadler, J. E. & Tulinsky, A. Structure of the Ser195Ala mutant of human alpha-thrombin complexed with fibrinopeptide A(7--16): evidence for residual catalytic activity. Acta. Crystallogr. D. Biol. Crystallogr. 56, 406–410 (2000).

20. Carter, P. & Wells, J. A. Dissecting the catalytic triad of a serine protease. Nature 332, 564–568 (1988).

21. Elshahawi, S. I., Shaaban, K. A., Kharel, M. K. & Thorson, J. S. A comprehensive review of glycosylated bacterial natural products. Chem. Soc. Rev. 44, 7591–7697 (2015).

22. Morar, M., Pengelly, K., Koteva, K., Wright, G. D. Mechanism and diversity of the erythromycin esterase family of enzymes. Biochemistry 51, 1740–1751 (2012).

23. Kim, D.W. & Cha, C-J. Antibiotic resistome from the One-Health perspective: understanding and controlling antimicrobial resistance transmission. Exp. Mol. Med. 53, 301 – 309 (2021).

24. Holm, L. & Laasko, L. M. Dali server update. Nucleic Acids Res. 44, W351–W355 (2016).

25. Chatonnet, A., Perochon, M., Velluet, E. & Marchot, P. The ESTHER database on alpha/beta hydrolase fold proteins - An overview of recent developments. Chem. Biol. Interact. 383, 110671 (2023).

26. Jansson, A., Niemi, J., Mäntsälä, P. & Schneider, G. Crystal structure of aclacinomycin methylesterase with bound product analogues: implications for anthracycline recognition and mechanism. J. Biol. Chem. 278, 39006–390013 (2003).

27. Rangel Pereira, P., Balan, A. & Hyvonen, M. Novel lipases isolated from metagenomic library [Unpublished]. Worldwide Protein Data Bank. PDB ID: 8PZG. PDB DOI: 10.2210/pdb8PZG/pdb (2024).

28. Vinces, T., et al. Monomeric esterase: Insights into cooperative behavior, hysteresis/allokairy. Biochemistry 63, 1178–1193 (2024).

29. Faulkner, N., et al. Characterization of an isobutylene epoxide hydrolase (IbcK) from the isobutylene-catabolizing bacterium Mycolicibacterium sp. ELW1. Appl. Environ. Microbiol. 91, e0039325 (2025).

30. Ruzzini, A. C., Bhowmik, S., Yam, K. C., Ghosh, S., Bolin, J. T. & Eltis, L. D. The lid domain of the MCP hydrolase DxnB2 contributes to the reactivity toward recalcitrant PCB metabolites. Biochemistry 52, 5685–5695 (2013).

31. Garcia-Viloca, M., Gao, J., Karplus, M. & Truhlar, D. G. How Enzymes Work: Analysis by Modern Rate Theory and Computer Simulations. Science 303, 186–195 (2004).

32. Davis, J. H., Rubin, A. J. & Sauer, R. T. Design, construction and characterization of a set of insulated bacterial promoters. Nucleic Acids Res. 39, 1131–1141 (2011).

33. Otwinowski, Z. & Minor, W. Processing of X-ray diffraction data collected in oscillation mode. Methods Enzymol. 276, 307–326 (1997).

34. Liebschner, D., et al. Macromolecular structure determination using X-rays, neutrons and electrons: recent developments in Phenix. Acta. Crystallogr. D. Struct. Biol. 75, 961–877 (2019).

35. Jumper, J., et al. Highly accurate protein structure prediction with AlphaFold. Nature 596, 583–590 (2021).

36. Emsley, P., Lohkamp, B., Scott, W. G. & Cowtan, K. Features and development of Coot. Acta. Crystallogr. D. Struct. Biol. 66, 486–501 (2010).

37. Transtrum, M. K., Hansen, L. D. & Quinn, C. Enzyme kinetics determined by single-injection isothermal titration calorimetry. Methods 76, 194–200 (2015).

38. Quinn, C. F. & Hansen, L. D. Enzyme Kinetics Determined by Single-Injection Isothermal Titration Calorimetry. Methods. Mol. Biol. 1964, 241–249 (2019).

39. Di Trani, J. M., Moitessier, N. & Mittermaier, A. K. Complete kinetic characterization of enzyme inhibition in a single isothermal titration calorimetric experiment. Anal. Chem. 90, 8430–8435 (2018).

40. Pei, J. & Grishnin, N. V. AL2CO: calculation of positional conservation in a protein sequence alignment. Bioinformatics 17, 700–712 (2001).

