## Supplemental Material for "Hydration and hydrolysis define antibiotic resistance conferred by macrolide esterases"

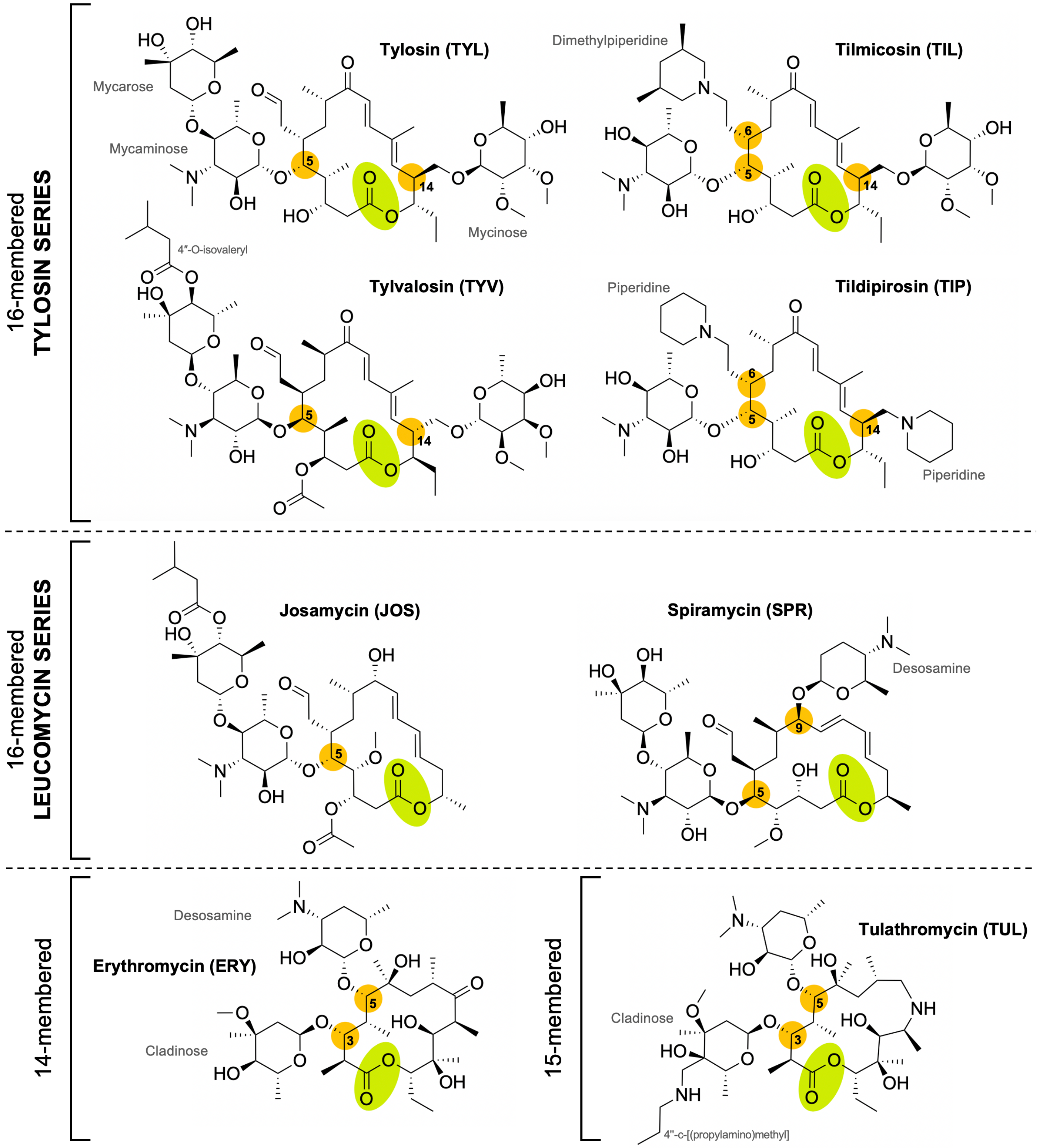


**Figure S1. Chemical structures of 14-, 15-, and 16-memebed macrolides tested for esterase activity.** Ring position substitutions have been highlighted in orange, ester bond highlighted in green.

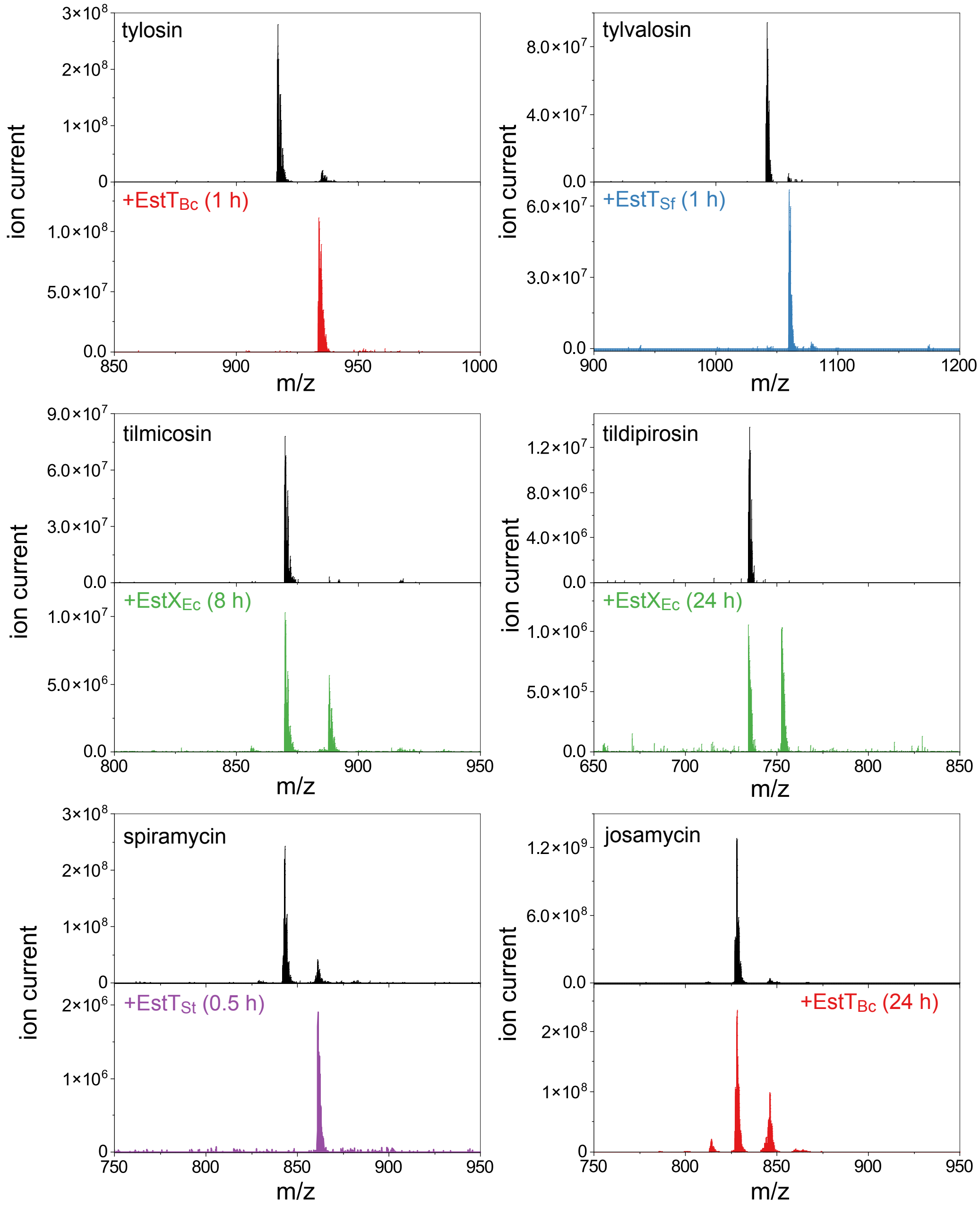


**Figure S2. Esterase-mediated macrolide hydrolysis.** Representative mass spectra demonstrating the hydrolysis of six macrolides by EstT and EstX homologues. In each panel, the top spectrum shows the putative macrolide substrate (black), the bottom panel shows hydrolysis by one of the known or putative macrolide esterases and the time at which the reaction was quenched with acetic acid

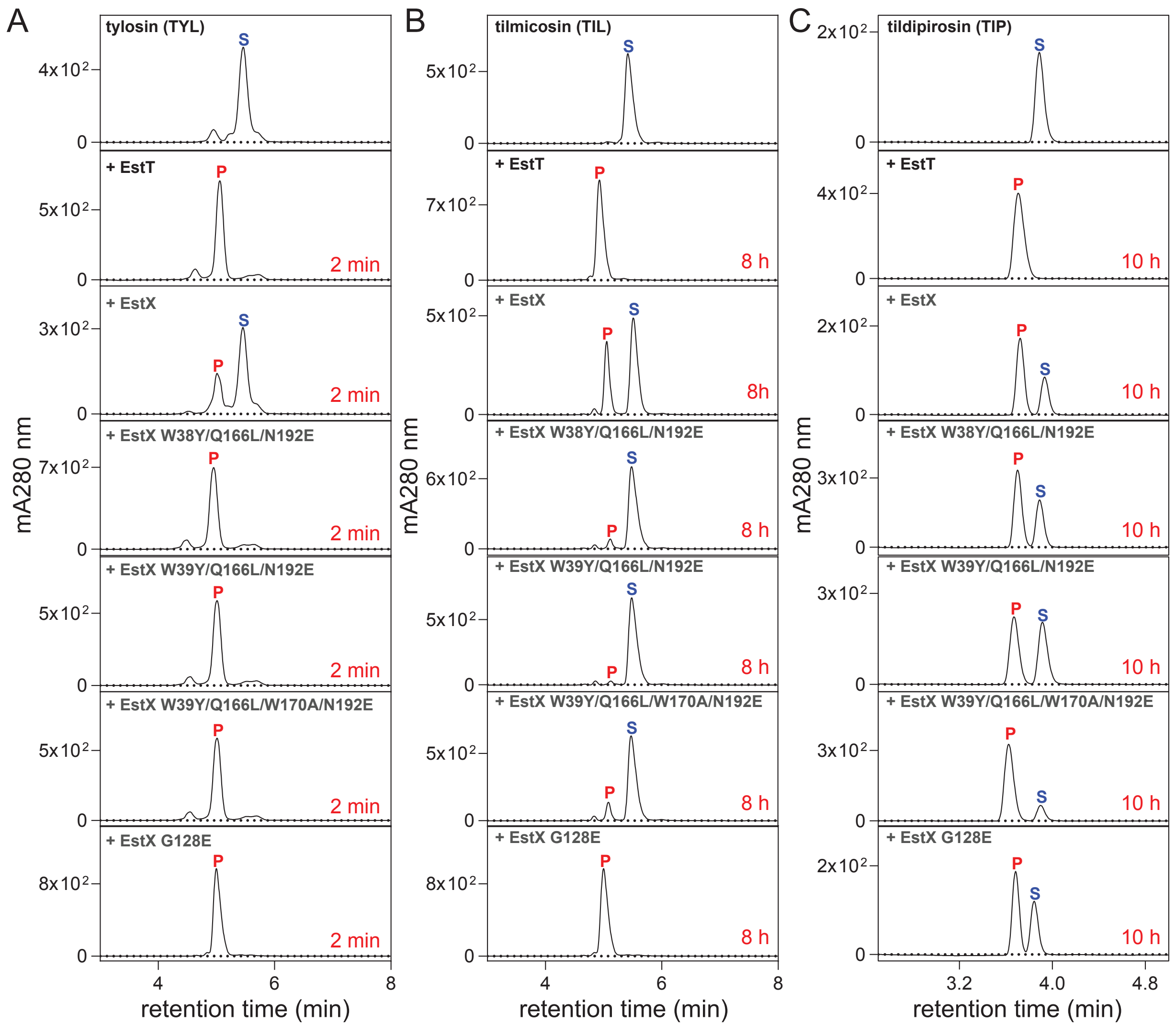


**Figure S3. Qualitative assessment of EstX$_{EC}$ variant activities.** HPLC/UV chromatograms of quenched reactions of each EstX variant (1 mM) with 200 mM **a.** TYL after 2 min, **b.** TIL after 8 h and **c.** TIP after 10 h. In each series of a panels, a quenched substrate control is show at the top followed by reactions catalysed by wildtype enzymes EstT$_{Sf}$ and EstX$_{Ec}$ for comparison.

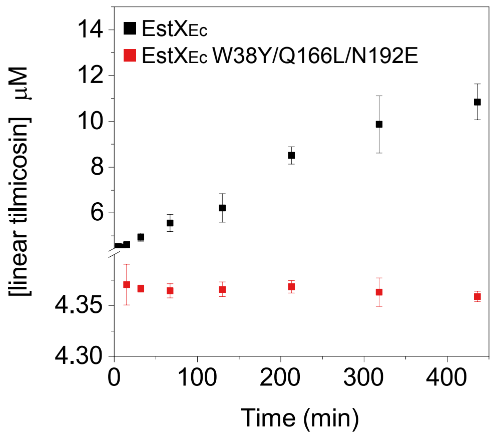


**Figure S4. Slow and abrogated EstX$_{Ec}$ turnover of tilmicosin.** Time-dependent quantification of linear TIL observed upon reaction with 510 nM EstX$_{Ec}$ (black) and EstX$_{Ec}$ W38Y/Q166L/N192E (red). No reaction between the latter and TIL was observed, indicated by no change in the concentration of linear TIL present in all samples: ~4.4 mM or 1.1% linear TIL is present in the commercial stock

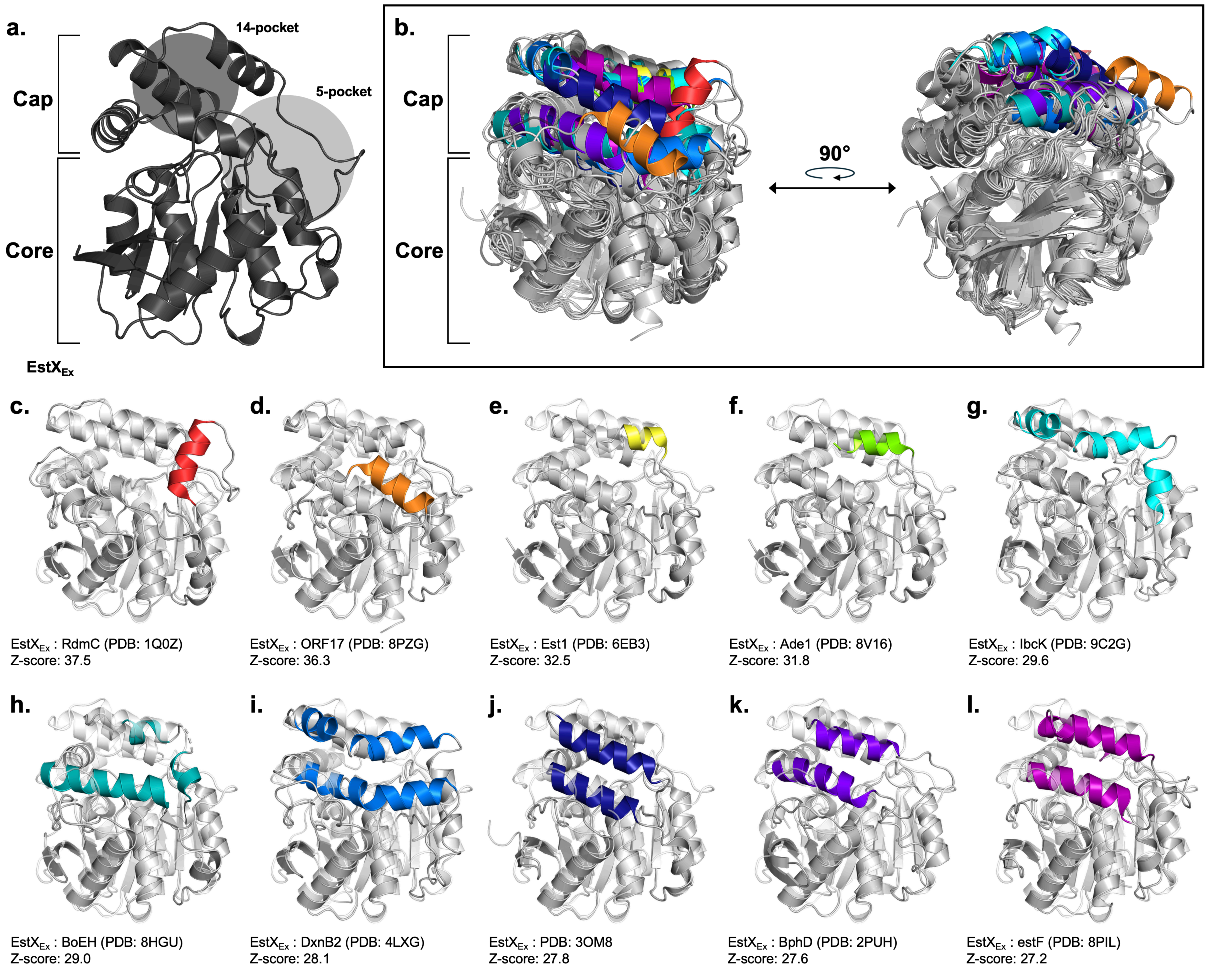


**Figure S5.** EstX$_{Ec}$ crystal structure **a.** compared to **b.** top ten closest Est enzyme structural homologs obtained via Dali server full PDB search. **c.-l.** Ten closest (by Dali Z-score) non-Est-type α/β-hydrolase structural homologs. Key structural differences are highlighted in colours other than grey.

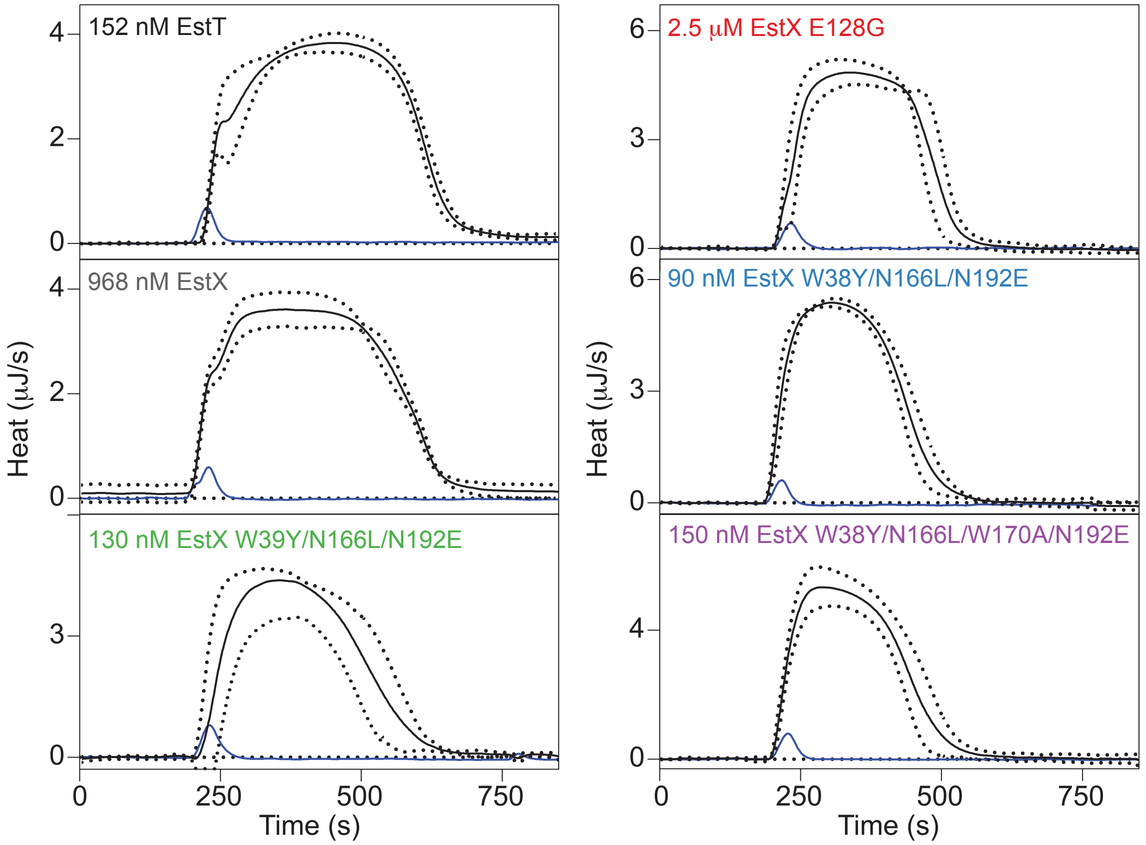


**Figure S6. Thermograms of single-injection ITC measurements of enzymatic reactions.** All reactions were performed at room temperature in 100 mM KPi, pH 7.0. The enzymes and concentrations are labeled at the top of each individual panel. The average thermal inputs from three reactions are shown as solid black lines, dotted lines show standard deviations between the reactions, and representative control injections of buffer into enzyme are drawn in dark blue.

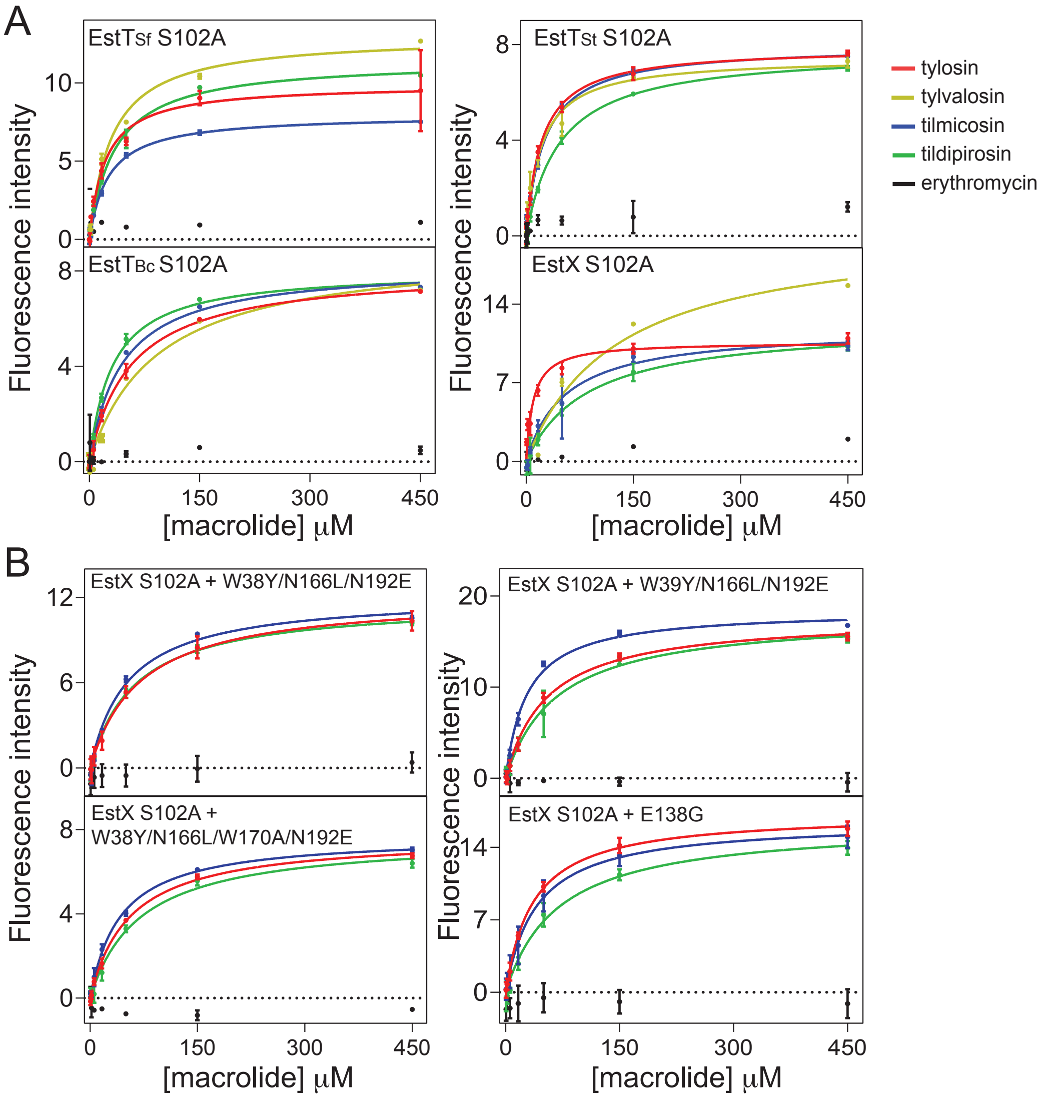


**Figure S7. Equilibrium binding experiments. a.** Binding curves showing titrations of catalytically inactive S102A EstT$_{Sf}$, EstT$_{St}$, EstT$_{Bc}$ and **b.** EstX$_{Ec}$ variants with TYL (red), TYV (yellow), TIL (blue), TIP (green) and ERY (black) monitored by changes in intrinsic tryptophan fluorescence.

**Supplemental Tables**

**Table S1.** Pairwise sequence and structural comparison between Est enzymes

| | $EstT_{Sf}$ | $EstT_{St}$ | $EstT_{Bc}$ | $EstX_{Ec}$ |
|---|---|---|---|---|
| $EstT_{Sf}$ | | 66 | 51 | 44 |
| $EstT_{St}$ | 0.8 | | 51 | 44 |
| $EstT_{Bc}$ | 1.4 | 1.1 | | 51 |
| $EstX_{Ec}$ | 1.3 | 1.1 | 1.2 | |

Values in blue boxes provide %identity; values in orange boxes give r.m.s.d. deviation in Å. All values were calculated using Dali [#].

**Table S2.** MICs[a] (μg/mL) of macrolides in LB + 1 mM EDTA[b]

| | TYL | TVY | TIL | TIP | SPM | ERY |
|---|---|---|---|---|---|---|
| *estT*$_{Sf}$ | 256-512 | 32 | 128 | 64 | 32 | 8 |
| *estT*$_{St}$ | 256 | 16-32 | 64 | 8 | 64 | 8 |
| *estT*$_{Bc}$ | 256-512 | 16-32 | 16 | ≤8 | 32 | 8 |
| *estX*$_{Ec}$ | 256 | 8-16 | 8 | ≤8 | 16 | 8 |
| *ereA* | 64 | 8 | 4 | ≤8 | 16 | 256-512 |
| *DH5α* | 64 | 16 | 4 | ≤8 | 8 | 8 |

*a* –MICs performed by 3 individuals in triplicate, ranges are presented if variation was observed
*b* – The addition of significantly impacts the MICs of macrolides (see Table S3).

.

**Table S3.** Macrolide esterase X-ray crystal diffraction data and refinement statistics

| | | $\mathrm{EstT_{Sf}\text{-}S_{cat}}$→A | $\mathrm{EstT_{St}\text{-}S_{cat}}$→A | $\mathrm{EstT_{Bc}\text{-}S_{cat}}$→A: L-TYV | $\mathrm{EstX_{Ec}\text{-}S_{cat}}$→A: L-TYL | $\mathrm{EstX_{Ec}\text{-}S_{cat}}$→A: L-TYV |
|---|---|---|---|---|---|---|
| *Data Collection* | | | | | | |
| Wavelength (Å) | | 1.03 | 1.18 | 1.03 | 1.18 | 1.18 |
| Resolution Range (Å) | | 29.44 - 3.19 (3.34 - 3.19) | 29.09 – 2.18 (2.23 – 2.18) | 29.51 – 1.7 (1.73 – 1.7) | 29.02 – 1.65 (1.69 – 1.65) | 29.04 – 1.99 (2.04 – 1.99) |
| Space Group | | $P\ 2_1\ 2_1\ 2_1$ | $P\ 2_1\ 2_1\ 2_1$ | $P\ 4_1\ 2_1\ 2$ | $P\ 1\ 2_1\ 1$ | $P\ 1\ 2_1\ 1$ |
| Unit Cell | a, b, c (Å) | 49.9, 82.3, 392.6 | 46.8, 74.3, 77.0 | 72.5, 72.5, 129.2 | 44.6, 72.7, 82.1 | 44.9, 72.7, 82.3 |
| | a, b, g (°) | 90, 90, 90 | 90, 90, 90 | 90, 90, 90 | 90, 95.7, 90 | 90, 95.2, 90 |
| Molecule copies in ASU | | 5 | 1 | 1 | 2 | 2 |
| Total reflections | | 210,342 (18,752) | 136,506 (2,226) | 949,348 (30,740) | 369,642 (1,124) | 185,505 (7,035) |
| Unique reflections | | 44,037 (4,512) | 26,538 (702) | 72,097 (2,899) | 112,953 (520) | 66,069 (3,419) |
| Completeness (%) | | 83.6 (54.1) | 74.9 (26.9) | 92.6 (53.0) | 95.8 (60.9) | 92.06 (58.4) |
| Mean I/sigma(I) | | 8.00 (2.73 | 8.37 (1.71) | 17.93 (2.27) | 10.49 (2.65) | 6.42 (2.40) |
| Wilson B-factor (Å$^2$) | | 59.01 | 26.30 | 12.06 | 11.61 | 22.10 |
| R-merge | | 0.13 (0.42) | 0.17 (0.71) | 0.1142 (0.8801) | 0.08 (0.29) | 0.11 (0.31) |
| CC1/2 | | 0.99 (0.694 | 0.986 (0.373) | 0.999 (0.72) | 0.995 (0.702) | 0.986 (0.700) |
| CC* | | 0.99 (0.91) | 0.996 (0.737) | 1 (0.915) | 0.999 (0.908) | 0.996 (0.907) |
| Multiplicity | | 4.8 (4.2) | 5.1 (3.2) | 13.2 (10.6) | 3.3 (2.2) | 2.8 (2.1) |
| *Refinement Statistics* | | | | | | |
| No. of reflections | | 23,456 (1873) | 12,998 (486) | 38,509 (1,508) | 60,035 (2,703) | 185,505 (7,035) |
| No. used for $R_{free}$ | | 1180 (106) | 1314 (50) | 2,000 (78) | 2,000 (90) | 66,069 (3,419) |
| $R_{work}/R_{free}$ | | 0.2649/0.2965 (0.3140/0.3916) | 0.1838/0.2375 (0.1972/0.3627) | 0.1777/0.2011 (0.2635/0.3554) | 0.1642/0.1988 (0.2112/0.2641) | 0.1637/0.2110 (0.1559/0.2196_ |
| Average B-factor (Å$^2$) | | 55.95 | 28.36 | 15.74 | 13.83 | 22.98 |
| Clashscore | | 1.70 | 4.80 | 0 | 4.34 | 3.59 |
| No. of atoms | Non-hydrogens | 9,647 | 2,132 | 2,356 | 4,765 | 4,583 |
| | Macromolecule | 9,633 | 2,037 | 2,058 | 4,311 | 4,301 |
| | Ligand | 0 | 0 | 74 | 130 | 74 |
| | Solvent | 14 | 95 | 224 | 324 | 208 |
| Ramachandran | Favoured (%) | 96.06 | 96.47 | 96.99 | 97.12 | 96.54 |
| | Allowed (%) | 3.86 | 3.14 | 3.01 | 2.52 | 3.46 |
| | Outliers (%) | 0.08 | 0.39 | 0 | 0.36 | 0.00 |
| Rotamer Outliers (%) | | 0.11 | 0.95 | 0 | 0.45 | 0.44 |

**Table S4.** Atom contacts formed between linearized macrolide, waters, and Est protein

| Protein | $EstT_{St}$ | | | $EstT_{Bc}$ : L-TLV | | | $EstX_{Ec}$ : L-TYL | | | $EstX_{Ec}$ : L-TLV | | |
|---|---|---|---|---|---|---|---|---|---|---|---|---|
| | Atom Contacts | | Å* | Atom Contacts | | Å | Atom Contacts | | Å | Atom Contacts | | Å |
| **5-pocket** | Water[i] | $O_{H2O}$ --- $O_{P129}$ | 2.8 | Water | $O_{H2O}$ --- $OG_{S128}$ | 2.7 | Water | $O_{H2O}$ --- $OE1_{E128}$ | 2.6 | Water | $O_{H2O}$ --- $OE1_{E128}$ | 3.1 |
| | | $O_{H2O}$ --- $N_{G131}$ | 3.1 | | $O_{H2O}$ --- $O_{V129}$ | 2.8 | | $O_{H2O}$ --- $O_{R129}$ | 2.7 | | $O_{H2O}$ --- $O_{R129}$ | 2.9 |
| | | | | | $O_{H2O}$ --- $O_{G131}$ | 3.1 | | $O_{H2O}$ --- $NE_{R129}$ | 2.9 | | $O_{H2O}$ --- $NH2_{R129}$ | 2.7 |
| | | | | | $O_{H2O}$ --- $OD2_{D133}$ | 2.8 | | $O_{H2O}$ --- $NH2_{R129}$ | 3.0 | | $O_{H2O}$ --- $O_{D132}$ | 3.3 |
| | | | | | $O_{H2O}$ --- $OD2_{D133}$ | 2.9 | | $O_{H2O}$ --- $OD1_{D132}$ | 3.4 | | $O_{H2O}$ --- $OD1_{D132}$ | 3.0 |
| | | | | | $O_{H2O}$ --- $OG1_{T136}$ | 2.7 | | $O_{H2O}$ --- $N_{A133}$ | 2.9 | | $O_{H2O}$ --- $N_{A133}$ | 3.0 |
| | | | | β-1- Mcn[ii] | $O_{H2O}$ --- $O34_{TVL}$ | 3.4 | | $O_{H2O}$ --- $N_{D134}$ | 2.9 | α-1,4-Mcr | $O_{H2O}$ --- $O48_{TVL}$ | 3.0 |
| | | | | | $O_{H2O}$ --- $O37_{TVL}$ | 3.0 | | $O_{H2O}$ --- $ND1_{H240}$ | 2.8 | | $O_{H2O}$ --- $O50_{TVL}$ | 3.4 |
| | | | | | $O_{H2O}$ --- $N39_{TVL}$ | 2.8 | β-1- Mcn | $O_{H2O}$ --- $O48_{TYL}$ | 2.7 | | $O37_{TYL}$ --- $OE2_{E128}$ | 2.8 |
| | | | | | $O_{H2O}$ --- $O43_{TVL}$ | 3.0 | | $O37_{TYL}$ --- $OE2_{E128}$ | 2.7 | | | |
| | | | | α-1,4-Mcr[iii] | $O_{H2O}$ --- $O48_{TVL}$ | 2.8 | | | | | | |
| | | | | | $O_{H2O}$ --- $O48_{TVL}$ | 3.0 | | | | | | |
| | | | | | $O_{H2O}$ --- $O50_{TVL}$ | 3.2 | | | | | | |
| | | | | | $O56_{TVL}$ --- $N_{G131}$ | 3.0 | | | | | | |
| **Ring-pocket** | Water | $O_{H2O}$ --- $N_{A32}$ | 2.8 | Lactone Ring (linear) | $O72_{TLV}$ --- $N_{A32}$ | 3.0 | Water | $O_{H2O}$ --- $OG_{S34}$ | 2.8 | Water | $O_{H2O}$ --- $O_{A32}$ | 2.9 |
| | | $O_{H2O}$ --- $OH_{Y38}$ | 2.7 | | $O72_{TLV}$ --- $N_{L103}$ | 2.9 | | $O_{H2O}$ --- $O_{S-CAT\rightarrow A}$ | 2.7 | | $O_{H2O}$ --- $NE2_{H261}$ | 2.8 |
| | | $O_{H2O}$ --- $N_{L103}$ | 2.9 | | $O73_{TLV}$ --- $NE2_{H263}$ | 2.8 | | $O_{H2O}$ --- $NE2_{H148}$ | 2.8 | | $O_{H2O}$ --- $ND2_{N173}$ | 2.7 |
| | | $O_{H2O}$ --- $NE2_{H260}$ | 2.8 | | | | | $O_{H2O}$ --- $ND2_{N173}$ | 2.8 | Lactone Ring (linear) | $O_{H2O}$ --- $O04_{TVL}$ | 2.8 |
| | | $O_{H2O}$ --- $O_{H260}$ | 2.7 | | | | | $O_{H2O}$ --- $NE2_{H261}$ | 2.9 | | $O_{H2O}$ --- $O72_{TVL}$ | 2.5 |
| | | | | | | | Ring | $O_{H2O}$ --- $O60_{TYL}$ | 2.8 | | $O_{H2O}$ --- $O72_{TVL}$ | 2.6 |
| | | | | | | | | $O_{H2O}$ --- $O63_{TYL}$ | 2.7 | | $O73_{TYL}$ --- $N_{A32}$ | 2.9 |
| | | | | | | | | $O_{H2O}$ --- $O63_{TYL}$ | 2.7 | | $O73_{TYL}$ --- $N_{L103}$ | 2.8 |
| | | | | | | | | $O_{H2O}$ --- $O65_{TYL}$ | 3.1 | | | |
| | | | | | | | | $O64_{TYL}$ --- $N_{A32}$ | 2.8 | | | |
| | | | | | | | | $O64_{TYL}$ --- $N_{L103}$ | 2.9 | | | |
| **14-pocket** | Water | $O_{H2O}$ --- $O_{D180}$ | 3.0 | Water | $O_{H2O}$ --- $OG_{S172}$ | 2.8 | Water | $O_{H2O}$ --- $N_{G63}$ | 3.0 | water | $O_{H2O}$ --- $O_{M33}$ | 2.6 |
| | | $O_{H2O}$ --- $OD2_{D180}$ | 3.3 | | $O_{H2O}$ --- $OG_{S172}$ | 2.8 | | $O_{H2O}$ --- $ND2_{N192}$ | 3.0 | | $O_{H2O}$ --- $N_{S35}$ | 3.0 |
| | | $O_{H2O}$ --- $O_{L262}$ | 3.4 | | $O_{H2O}$ --- $OG1_{T33}$ | 2.9 | | $O_{H2O}$ --- $O_{M33}$ | 2.8 | | $O_{H2O}$ --- $ND2_{N192}$ | 3.3 |
| | | $O_{H2O}$ --- $N_{L262}$ | 2.9 | | $O_{H2O}$ --- $NE2_{H150}$ | 2.8 | | $O_{H2O}$ --- $O_{A32}$ | 2.9 | | $O_{H2O}$ --- $OG1_{T196}$ | 2.9 |
| | | $O_{H2O}$ --- $OG1_{T264}$ | 2.8 | β-1-Myc[iv] | $O16_{TVL}$ --- $O_{L168}$ | 2.9 | β-1-Myc | $O_{H2O}$ --- $O16_{TYL}$ | 3.4 | β-1-Myc | $O_{H2O}$ --- $O16_{TVL}$ | 3.4 |
| | | $O_{H2O}$ --- $N_{T264}$ | 3.1 | | $O_{H2O}$ --- $O19_{TVL}$ | 3.1 | | $O_{H2O}$ --- $O07_{TYL}$ | 3.0 | | $O_{H2O}$ --- $O07_{TVL}$ | 3.3 |
| | | $O_{H2O}$ --- $N_{L262}$ | 2.9 | | $O_{H2O}$ --- $O10_{TVL}$ | 2.8 | | $O60_{TYL}$ --- $N_{L103}$ | 3.4 | | $O16_{TYL}$ --- $OE1_{Q166}$ | 3.0 |
| | | | | | | | | $O16_{TYL}$ --- $OE1_{Q166}$ | 2.9 | | $O16_{TYL}$ --- $ND2_{N192}$ | 3.1 |
| | | | | | | | | $O16_{TYL}$ --- $ND2_{N192}$ | 2.9 | | | |
| | | | | | | | | $O_{H2O}$ --- $O04_{TYL}$ | 2.7 | | | |

* Atomic Contact Distance in Å
[i] Waters in contact with esterase
[ii] β-1-mycaminose (Mcn) contacts with water and esters
[iii] α-1,4-mycarose (Mcr) contacts with water and esters
[iv] β-1-mycinose (Myc) contacts with water and esters

**Table S5.** Dissociation constants ($K_D$, mM) measured for catalytically inactive esterases and macrolides

| $S_{cat}$→A variant | TYL | TIL | TIP |
|---|---|---|---|
| EstX | 10 ± 2 | 55 ± 14 | 86 ± 15 |
| E128G | 36 ± 3 | 40 ± 7 | 69 ± 11 |
| W38Y/Q166L/N192E | 69 ±10 | 53 ± 7 | 63 ± 5 |
| W39Y/Q166L/N192E | 54 ± 5 | 28 ± 2 | 68 ± 11 |
| W38Y/Q166L/W170A/N192E | 55 ± 4 | 42 ± 3 | 70 ± 9 |

ND: not determined

Some of this data is also shown in Table 3.

**Table S6.** Summary of relevant Dali Full PDB search results compared to macrolide esterases EstT$_{Sf}$, EstT$_{St}$, EstT$_{Bc}$, and EstX$_{Ec}$

| Protein | PDB | Function & Class | %ID [i] | | | | RMSD (Å) | | | | z-score | | | | |
|---|---|---|---|---|---|---|---|---|---|---|---|---|---|---|---|
| | | | EstT$_{Sf}$ | EstT$_{St}$ | EstT$_{Bc}$ | EstX$_{Ec}$ | EstT$_{Sf}$ | EstT$_{St}$ | EstT$_{Bc}$ | EstX$_{Ec}$ | EstT$_{Sf}$ | EstT$_{St}$ | EstT$_{Bc}$ | EstX$_{Ec}$ | Mean: |
| RdmC | 1Q0Z | Methylesterase [A] | 28 | 30 | 31 | 34 | 2.1 | 1.9 | 2.0 | 2.0 | 36.5 | 35.6 | 35.1 | 37.5 | 36.1 |
| ORF17 | 8PZG | Lipase [A] | 29 | 28 | 29 | 29 | 2.0 | 1.9 | 1.8 | 1.9 | 35.8 | 34.1 | 35.2 | 36.3 | 35.6 |
| Est1 | 6EB3 | Esterase [A] | 28 | 31 | 30 | 25 | 2.1 | 2.0 | 1.9 | 2.0 | 32.1 | 31.5 | 31.6 | 32.5 | 32.0 |
| Ade1 | 8V16 | Esterase [A] | 28 | 30 | 30 | 25 | 2.1 | 2.1 | 2.1 | 2.2 | 31.6 | 31.8 | 31.4 | 31.8 | 31.7 |
| IbcK | 9C2G | Epoxide hydrolase [A] | 19 | 20 | 18 | 19 | 2.5 | 2.3 | 2.6 | 2.5 | 29.1 | 29.1 | 28.4 | 29.6 | 29.1 |
| BoEH | 8HGU | Epoxide hydrolase [A] | 17 | 21 | 18 | 20 | 2.6 | 2.1 | 2.4 | 2.4 | 28.8 | 28.3 | 28.9 | 29.0 | 28.8 |
| DxnB2 | 4LXG | C-C hydrolase [A] | ••• [iii] | 16** | 19 | 16 | ••• | 2.7** | 2.6 | 2.6 | ••• | 28.0** | 28.2 | 29.0 | 28.6 |
| --- [ii] | 3OM8 | Lactonase [A*] | 21 | 22 | 24 | 21 | 2.6 | 2.5 | 2.7 | 2.8 | 28.3 | 28.8 | 27.4 | 28.1 | 28.2 |
| BphD | 2PUH | C-C hydrolase [A] | 19 | 18 | 23 | 20 | 2.5 | 2.7 | 2.4 | 2.6 | 28.4 | 28.4 | 27.4 | 27.8 | 28.0 |
| estF | 8PI1 | Arylesterase [A] | 17 | 18 | 21 | 19 | 2.6 | 2.8 | 2.6 | 2.7 | 28.0 | 27.4 | 28.0 | 27.6 | 27.8 |
| PcaD | 2XUA | Enol-lactonase [A] | 22 | 21 | 20 | 19 | 2.4 | 2.6 | 2.8 | 3.0 | 27.6 | 28.4 | 27.3 | 27.2 | 27.6 |
| --- | 3FOB | Bromoperoxidase [B] | 20 | 20 | 21 | 21 | 2.7 | 2.8 | 2.5 | 2.8 | 27.5 | 27.1 | 27.6 | 28.1 | 27.6 |
| PhaZ | 8YNW | Depolymerase [A] | 20 | 23 | 26 | 21 | 2.4 | 2.3 | 2.2 | 2.6 | 27.1 | 28.3 | 27.1 | 26.9 | 27.3 |
| CpoF | 1A8S | Chloroperoxidase [c] | 18 | 21 | 21 | 20 | 2.6 | 2.9 | 2.7 | 2.8 | 26.8 | 26.7 | 27.2 | 27.1 | 27.0 |
| Est816 | 5EGN | Lactonase [A] | 21 | 22 | 22 | 24 | 2.9 | 3.3 | 3.1 | 3.0 | 27.1 | 26.8 | 26.4 | 27.0 | 26.8 |
| MGS-M2 | 4Q3L | Carboxylesterase [A] | 19 | 19 | 22 | 20 | 3.0 | 3.1 | 2.7 | 3.1 | 27.1 | 27.1 | 26.3 | 26.6 | 26.8 |
| BpoC | 3E3A | Bromoperoxidase [B*] | 15 | 18 | 21 | 18 | 2.9 | 2.6 | 2.6 | 2.9 | 26.9 | 27.4 | 26.6 | 26.3 | 26.8 |
| TtEst | 4UHD | Esterase [A] | 19 | 19 | 24 | 21 | 2.7 | 2.7 | 3.3 | 2.9 | 27.1 | 27.5 | 25.6 | 26.1 | 26.6 |
| BPO-A2 | 4ITV | Bromoperoxidase [B] | 17 | 20 | 21 | 24 | 2.7 | 2.8 | 2.8 | 2.7 | 26.6 | 26.1 | 26.4 | 26.3 | 26.4 |
| MehpH | 8HGV | C-C hydrolase [A] | 16 | 19 | 23 | 21 | 2.5 | 2.5 | 2.5 | 2.7 | 27.3 | 25.7 | 26.2 | 26.5 | 26.4 |
| Lip1 | 4OPM | Lipase [A*] | 19 | 19 | 21 | 21 | 2.5 | 2.7 | 2.6 | 2.8 | 27.4 | 26.8 | 25.7 | 25.8 | 26.4 |
| Lip3 | 6I8W | Phospholipase [A] | 16 | 21 | 20 | 20 | 2.7 | 2.6 | 2.6 | 19 | 27.3 | 26.8 | 25.5 | 26.0 | 26.4 |
| B1EPH2 | 6UNW | Epoxide hydrolase [A] | 19 | 16 | 21 | 20 | 2.8 | 2.5 | 2.8 | 2.6 | 26.8 | 26.6 | 25.0 | 26.0 | 26.1 |
| EphA | 5CW2 | Epoxide hydrolase [A*] | 22 | 18 | 20 | 18 | 2.7 | 2.6 | 2.7 | 2.8 | 26.5 | 26.8 | 25.1 | 25.4 | 26.0 |
| Ephx2 | 5ALJ | Epoxide hydrolase [A] | 15 | 18** | 17 | 19*** | 2.5 | 2.4** | 2.7 | 2.9*** | 26.7 | 26.5** | 25.3 | 26.3*** | 26.0 |
| Fac-dex | 3B12 | Dehalogenase [A] | 13 | 19 | 18 | 17 | 2.8 | 2.8 | 3.1 | 3.0 | 26.6 | 26.5 | 24.7 | 26.3 | 26.0 |
| EH2 | 3KXP | Epoxide hydrolase [A] | 19 | 18 | 18 | 20 | 2.7 | 2.6 | 2.6 | 2.9 | 25.9 | 26.2 | 25.9 | 25.5 | 25.9 |
| --- | 9C9F | Dehalogenase [A] | 14 | 19 | 15 | 15 | 3.9 | 2.8 | 3.0 | 3.1 | 26.2 | 25.4 | 25.2 | 25.9 | 25.7 |
| --- | 1HL7 | γ-lactamase [B] | 19 | 20 | 20 | 20 | 2.7 | 2.5 | 2.7 | 2.7 | 26.0 | 25.5 | 25.6 | 26.0 | 25.7 |
| DmrA | 4MJ3 | Dehalogenase [A] | 14 | 13 | 14 | 16 | 2.6 | 2.6 | 2.4 | 2.5 | 25.6 | 25.1 | 25.3 | 25.9 | 25.5 |
| MhpC | 9IJK | C-C hydrolase [A] | 18 | 16 | 20 | 21 | 2.6 | 2.8 | 2.7 | 2.7 | 26.1 | 24.9 | 24.4 | 25.1 | 25.1 |
| VACVase | 2OCK | Ser hydrolase [A] | 21 | 19 | 20 | 17 | 2.8 | 2.8 | 2.7 | 2.7 | 25.2 | 24.7 | 24.7 | 24.7 | 24.8 |
| PIP F1 | 1MTZ | Pro iminopeptidase [A] | 17 | 17 | 21 | 16 | 3.0 | 3.1 | 2.9 | 3.2 | 25.4 | 24.5 | 23.9 | 24.8 | 24.7 |
| --- | 5JYC | Hydrolase [A*] | 13 | 13** | 13 | 14*** | 2.7 | 2.8 | 3.0 | 2.9*** | 25.0 | 25.1 | 23.4 | 25.0*** | 24.5 |

* Putative

[A] Hydrolase
[B] Oxidoraductase
[C] Haloperoxidase

[i] %ID = percent identity of aligned amino acids
[ii] --- = Unnamed protein
[iii] ••• = No search result

** Related PDB matched to EstTSt: 4LYD, 4X6Y, 5TNK
*** Related PDB matched to EstX: 4Y2P, 3KD2

**Table S7.** Representative impact of EDTA on MICs (μg/mL) of macrolides

| | TYL | | ERY | |
|---|---|---|---|---|
| | No EDTA | 1 mM EDTA | No EDTA | 1 mM EDTA |
| $estT_{\mathrm{Sf}}$ | >1024 | 256-512 | 64 | 8 |
| $estT_{\mathrm{St}}$ | >1024 | 256 | 64 | 8 |
| $estT_{\mathrm{Bc}}$ | 512 | 256-512 | 64 | 8 |
| *estX* | >1024 | 256 | 64 | 8 |
| *ereA* | >1024 | 64 | 256-512 | >1024 |
| *DH5α* | 512 | 64 | 64 | 8 |

**Table S8.** MICs (μg/mL) of macrolides for *estX* and variants in LB and LB + 1 mM EDTA (LBE)

| ***E. coli* DH5α host** | **LB** | **LBE** |
|---|---|---|
| *estX* WT | 128 | 16 |
| E182G | 128 | 16 |
| W38Y/Q166L/N192E | 128 | <8 |
| W38Y/Q166L/W170A/N192E | 64 | <8 |
| W39Y/Q166L/N192E | 64 | 16 |

**Table S9.** Observed and theoretical ions for macrolides esterase-catalyzed hydrolysis[a]

| | **macrolide (substrate) ion [M+H]$^+$** | | **linear product ion [M+H]$^+$** | |
|---|---|---|---|---|
| | **theoretical** | **observed** | **theoretical** | **observed** |
| erythromycin | *733.9* | *735.1* | *only substrate observed*[b] | |
| tulathromycin | *806.0* | *807.1* | *only substrate observed*[b] | |
| spiramycin (SPM) | *843.5* | *844.0* | *861.5* | *862.0* |
| josamycin (JOS) | *828.0* | *829.1* | *846.0* | *847.1* |
| tylosin (TYL) | *916.2* | *917.2* | *934.2* | *935.2* |
| tylvalosin (TVL) | *1042.3* | *1043.3* | *1060.3* | *1061.3* |
| tilmicosin (TIL) | *869.1* | *870.2* | *887.1* | *888.2* |
| tildipirosin (TIP) | *734.0* | *735.1* | *752.0* | *753.1* |

*a* – values are provided to a 0.1 Da to match the accuracy of the instrument
*b* – no reactions observed after 24 h of incubation at RT in 100 mM KPi, pH 7.0
